# Identification and characterization of vasoactive intestinal peptide receptor antagonists with high-affinity and potent anti-leukemia activity

**DOI:** 10.1101/2024.11.08.622716

**Authors:** Yuou Wang, Anish Sen-Majumdar, Jian-Ming Li, Srijon Sarkar, Tenzin Passang, Yiwen Li, Jamie Cohen, Zihan Chen, Kiranj Chaudagar, Pankoj Kumar Das, Shuhua Wang, Nabute Bruk, Nikolaos Papadantonakis, Cynthia R. Giver, Edmund K. Waller

## Abstract

Vasoactive intestinal peptide (VIP) is a neuropeptide involved in tumor growth and immune modulating functions. Previous research indicated that a VIP antagonist (VIPhyb) enhances T-cell activation and induces T-cell-dependent anti-leukemic activity in mice. We created a combinatorial library of VIPhyb C-terminal sequence variations to develop a more potent VIP-receptor (VIP-R) antagonist, hypothesizing that specific amino acid substitutions would improve receptor binding and plasma stability. *In silico* screening analyses identified sequences with improved docking scores predicting increased binding affinity to human VIP receptors VPAC1 and VPAC2. Fifteen peptides were synthesized and tested for their ability to potentiate activation of purified mouse and human T cells and enhance T cell-dependent anti-leukemia responses in murine models of acute myeloid leukemia. Treating C57Bl/6 mice engrafted with a C1498 myeloid leukemia cell line with daily subcutaneous injections of VIP-R antagonist peptides induced T cell activation resulting in specific anti-leukemia responses. Strikingly, the predicted binding affinity of the VIP-R antagonists to VIP receptors correlated positively with their ability to augment mouse T-cell proliferation and anti-leukemia activity. ANT308 and ANT195 emerged as top candidates due to their high predicted VIP-R binding, low EC_50_ for *in vitro* T cell activation, and potent anti-leukemia activities. ANT308 decreased CREB phosphorylation, a downstream signaling pathway of the VIP receptor, and stimulated granzyme B and perforin expression in CD8+ T cells from AML patients. Combining *in silico* modeling, *in vitro* T cell activation properties, and *in vivo* anti-leukemia activity has identified promising VIP-R antagonist candidates for further development as novel immunotherapies for AML, especially for patients with relapsed disease.

## INTRODUCTION

Vasoactive intestinal peptide (VIP) is a 28 amino acid neuropeptide found in the brain and autonomic nervous system (1, 2), pancreas (3–5), and intestine (6–8). VIP is implicated in multiple physiologic processes, from gastrointestinal system homeostasis to neuronal development and modulation of immune response and inflammation (9). In the context of immune regulation, VIP secretion has been described as a characteristic of Th2 lymphocytes and non-lymphoid cells (9, 10) and binds with comparable affinity to VPAC1 and VPAC2, class B G-coupled protein receptors expressed on immune cells (11). VIP signaling activates cAMP-dependent pathways, promotes phosphorylation of cAMP response element binding protein (CREB), inhibits NF-κB activation, and exhibits anti-inflammatory and immunomodulatory capacities (12–15). Blockade of VIP signaling by peptides or antibodies leads to suppression of pancreatic (5, 16) and colon tumor growth(17, 18), and inhibits breast cancer cell migration (19), through modulation of tumor-intrinsic and T cell-dependent pathways. VIP over-expression in some solid-tumor cancers is mutually exclusive with expression of PD-L1, leading to the hypothesis that VIP receptor signaling is an alternative immune check-point pathway (20). However, the role of VIP expression on hematological malignancies has not been widely explored.

Acute myeloid leukemia (AML) is a common blood cancer in adults and is curable with standard chemotherapy only in a subset of patients. Allogeneic bone marrow transplant, a form of adoptive cellular immunotherapy, can improve outcomes, highlighting the role of the immune system in eradicating AML cells. Transcriptomic analysis showed 30% of acute myeloid leukemia patients overexpress VIP (21), suggesting that some AML tumors may utilize VIP as a mechanism to evade immune surveillance. Additionally, blood levels of VIP could serve as predictive biomarkers of clinical response and resistance (22). We have previously reported that inhibiting VIP receptor signaling with the VIP-R antagonist VIPhyb significantly enhances a T-cell-dependent, autologous anti-leukemia response in murine models of acute myeloid leukemia and T lymphoblastic leukemia (23) and augments anti-viral and anti-leukemia specific adaptive immunity in the treated animals (24–26).

The present study was undertaken to develop and evaluate higher-potency VIP-receptor antagonists for their ability to increase anti-leukemia specific immune activity. The starting point for developing VIP-receptor antagonists was VIPhyb, which contains the amino acid sequence of VIP modified by replacing the first six C-terminal amino acids with neurotensin sequences to create a competitive antagonist that binds to the VIP receptor but does not induce signaling (27–30). VIPhyb causes a half-maximal inhibition of VIP binding to VIP receptors on lymphocytes at 5 μM, and maximal inhibition of VIP-induced cAMP generation at 10 μM in experimental conditions (28). The low potency of VIPhyb has limited its suitability for clinical translation as a pharmacological VIP-receptor antagonist. Effective use of VIPhyb as an anti-cancer immune adjuvant in mouse leukemia models required pre-treatment of mice before leukemia inoculation (23), and less than 50% of treated animals achieved long-term leukemia-free survival. While concurrent inoculation of mice with tumors and treatment with VIPhyb demonstrated proof-of-principle evidence for the validity of VIP signaling as an immune check point, clinical translation of VIPhyb with this schedule of drug administration is not feasible (23).

To develop more potent VIP-R antagonists as potential anti-cancer immunotherapeutic, we performed rational drug design, identifying sequence variants of VIPhyb with enhanced potency and stability. We generated a combinatorial library of novel potential VIP-R antagonists (VIP-ANTs) based upon substitutions in the C-terminal peptide sequence of VIP that have been found in human cancer or the related peptide histidine isoleucine (PHI). We describe herein the selection of high-potency VIP-ANT from this combinatorial library based upon predicted binding affinities to human VPAC1 and VPAC2, confirm the increased potency of these VIP-R antagonist peptides in inducing autologous immune responses to myeloid malignancies in mice, and demonstrate *in vitro* effects on T cells purified from AML patients.

## RESULTS

### Generation of novel VIP receptors antagonists

The synthetic VIPhyb contains six N-terminal amino acids taken from neurotensin (residues 6-11) and the 22 C-terminal amino acids of VIP (Fig. 1A) (27–30). To develop VIP-R antagonists with enhanced predicted affinity, a library of 300 novel antagonists (VIP-ANTs) was created by adding single or multiple missense substitutions at positions 7-28 of VIPhyb (Fig. 1B). The specific base substitutions (T7A, D8V, Y10C, R12S, M17I, K20N, L23M, S25L) are naturally occurring cancer-associated mutations found in the Cancer Genome Atlas (TCGA) data portal (https://portal.gdc.cancer.gov). We hypothesized that these cancer-driven VIP variants might bind with higher affinity to VIP-R expressed on immune cells or be resistant to endopeptidases in blood, leading to improved pharmacokinetics and enhanced suppression of anti-cancer Th1/Tc1 T-cell activation in the tumor microenvironment (26). Also, since switching polar to hydrophobic residues like methionine in ligands have been observed to modulate receptor binding and signaling specificity (31–33), we tested N24M substitution in our screening (Fig. 1B). PHI shares 48% amino acid homology with VIP (14), PHI-derived D8S, N9D, I26L, L27I substitutions were included in our library as well (Fig. 1A). Lastly, substitutions of alanine or histidine were made at residues 1 to 28 in the amino acid sequence of VIP or VIPhyb (34–36)

**Figure 1.**
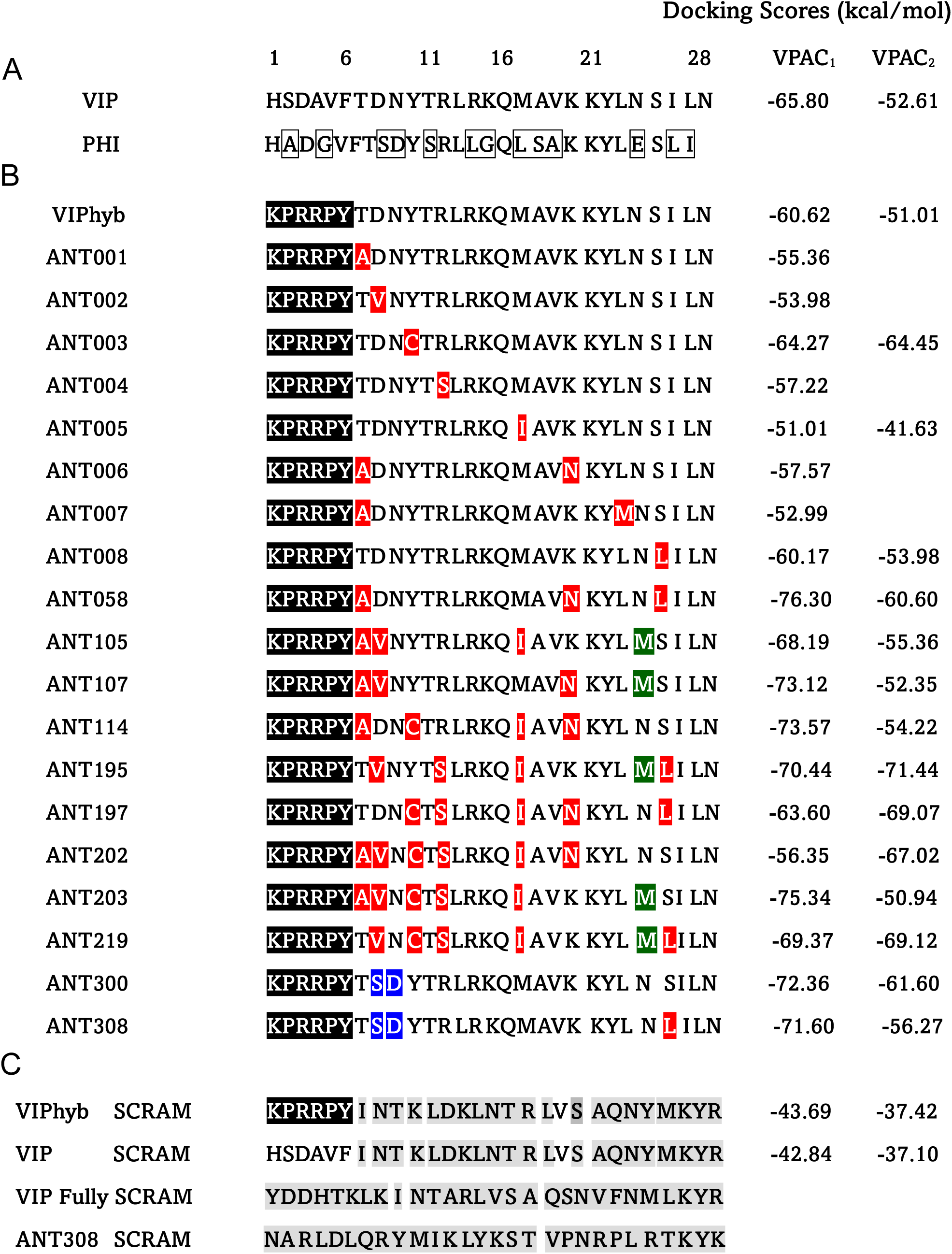
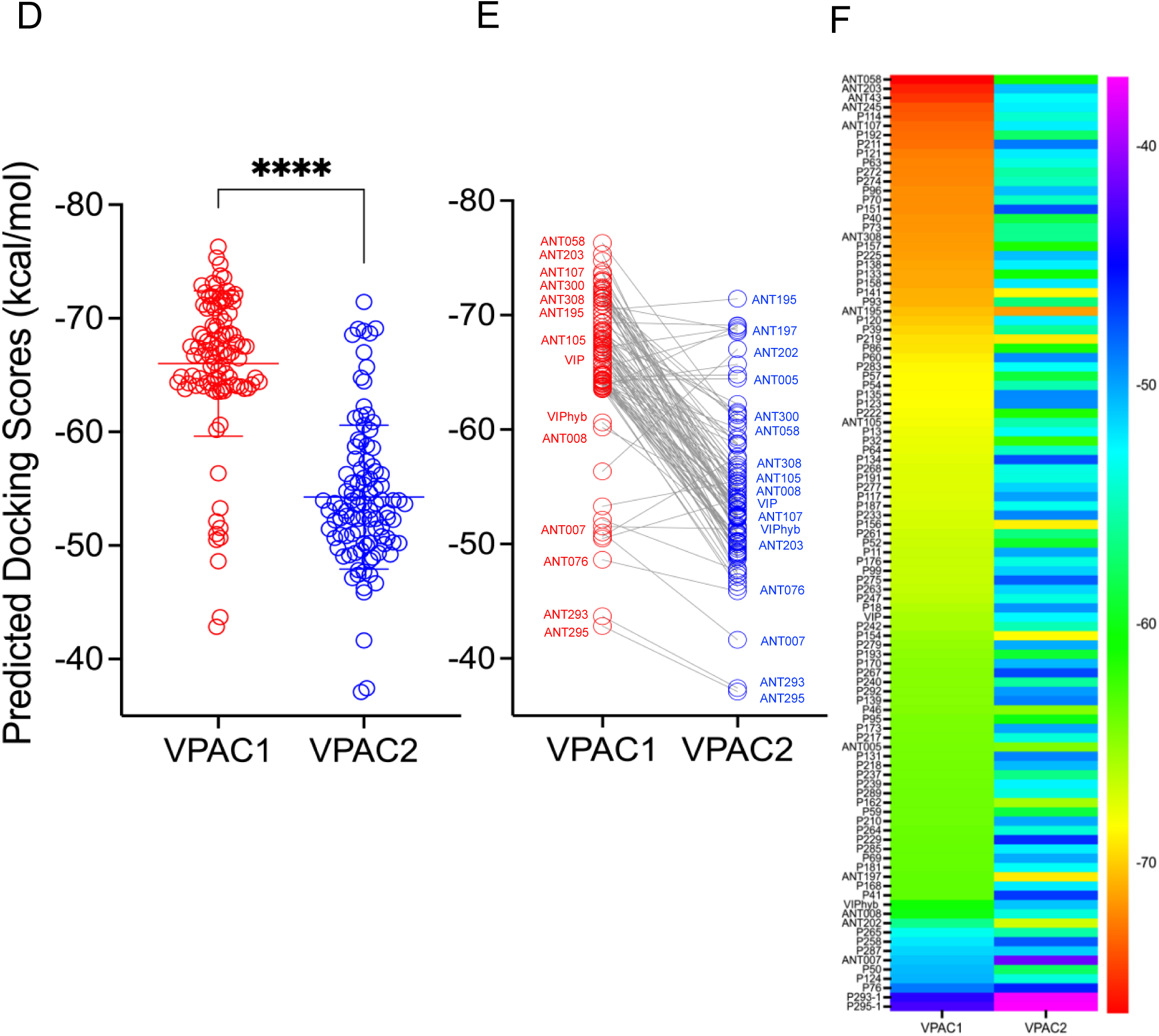
Sequence alignment of VIP, novel VIP antagonist peptides and docking scores. *(A)* Amino acid sequences of vasoactive intestinal peptide (VIP) and peptide histidine isoleucine (PHI). *(B)* Sequences of VIPhyb and selected peptides of the novel VIP antagonist (VIP-ANTs). The six amino acid fragments from neurotensin (residues 6– 11) are highlighted in *black*. C-terminal VIP mutations observed in human cancers are highlighted in *red*. Methionine substitutions are shaded in *green*. PHI homolog substitutions are shaded in *blue*. *In silico* docking scores predicting binding affinities for VPAC1 and VPAC2 are listed on the right. *(C)* Scrambled peptide sequences are shown. The divergence of amino acids from the parent sequence at specific positions is shaded in *grey*. *(D-F)* The top 100 VIP-ANTs’ docking scores to VPAC1 & 2 were graphed and compared as stacked scatter plots or heatmaps. ****P<0.0001, mean ± SD, unpaired t-test.

200 VIP-ANTs in the library were screened virtually for predicted binding to VPAC1. The top 100 variants with high predicted affinities to VPAC1 were subsequently analyzed for their docking scores to VPAC2 (Table 1 & Table 2). Docking scores for VIPhyb binding to VPAC1 and VPAC2 predict slightly lower affinities than VIP (Fig. 1A&B). As expected, VIPhyb SCRAM and VIP SCRAM, with random (scrambled) positions of amino acids at positions 7-28, had around 30% difference in docking scores predicting reduced affinity for both VPAC1&2, compared to the non-scrambled peptides, indicating the importance of the C-terminal for receptors binding (Fig. 1C). Noticeably, PHI homolog substitutions (D8S & N9D in ANT300 & ANT308) led to ∼20% change in docking scores predicting increased binding to VPAC1 (ANT300 & ANT308). Cancer-associated mutations (R12S, M17I & S25L in ANT195, ANT197 & ANT219) improved docking scores to VPAC2. Most VIP-ANTs have better docking scores for VPAC1 than VPAC2 (Fig. 1D&E); VPAC1 docking cannot predict VPAC2 docking (Fig. 1F).

**Table 1.** Summary of *in silico* screening results of novel VIP-ANTs. Scores for 100 peptide VIP-receptor antagonist sequences were categorized according to the predicted free energy of binding to VPAC1 by Creative Biolabs. The predicted binding affinity to VPAC2 was determined for the 100 peptides with the highest affinities to VPAC1. The categorization for the predicted affinity of VIP to VPAC1 and VPAC2 is shown in *bold*. Peptides with scrambled amino acid sequences had the lowest predicted binding affinity (> −50 kcal/mol).

| Range of Binding Score<br>(kcal/mol) | 200 Peptides<br>Screened for Binding<br>to VPAC1 | 100 Peptides Screened for<br>Binding to VPAC2 |
| --- | --- | --- |
| -70.1 to -76.3 | 26 peptides | 1 peptide |
| -65.1 to -70.0 | 35 peptides<br>**including VIP** | 7 peptides |
| -60.1 to -65.0 | 29 peptides | 9 peptides |
| -55.1 to -60.0 | 1 peptide | 18 peptides |
| -50.1 to -55.0 | 6 peptides | 44 peptides<br>**including VIP** |
| > -50.0 | 3 peptides including 2<br>scrambled ones | 21 peptides including 2 scrambled<br>ones |

**Table 2.** List of *in silico* docking scores of VIP-ANTs to Human VPAC1 & VPAC2. . Peptides are ordered based on their predicted affinities to VPAC1. VIP, VIPhyb, and ANT308 are shown as **<u>bold and underlined</u>**.

| VIP-ANTs | VPAC1 | VPAC2 | VIP-ANTs | VPAC1 | VPAC2 | VIP-ANTs | VPAC1 | VPAC2 |
| --- | --- | --- | --- | --- | --- | --- | --- | --- |
| ANT058 | -76.3 | -60.602 | ANT135 | -68.54 | -49.207 | ANT046 | -64.63 | -64.78 |
| ANT203 | -75.34 | -50.942 | ANT123 | -68.49 | -49.326 | ANT095 | -64.39 | -60.16 |
| ANT043 | -74.75 | -52.943 | ANT222 | -68.32 | -61.536 | ANT173 | -64.36 | -50.13 |
| ANT245 | -73.77 | -52.399 | ANT105 | -68.19 | -55.355 | ANT217 | -64.33 | -53.967 |
| ANT114 | -73.57 | -54.216 | ANT013 | -67.99 | -53.095 | ANT005 | -64.27 | -64.452 |
| ANT107 | -73.12 | -52.346 | ANT032 | -67.89 | -62.235 | ANT131 | -64.26 | -49.039 |
| ANT192 | -73 | -57.428 | ANT064 | -67.79 | -54.6 | ANT218 | -64.18 | -50.675 |
| ANT211 | -72.91 | -48.713 | ANT134 | -67.67 | -47.426 | ANT237 | -64.15 | 56.531 |
| ANT121 | -72.43 | -52.297 | ANT268 | -67.52 | -53.564 | ANT239 | -64.05 | -52.511 |
| ANT063 | -72.3 | -53.72 | ANT191 | -67.51 | -53.744 | ANT289 | -64.03 | -54.017 |
| ANT272 | -72.26 | -55.523 | ANT277 | -67.51 | -51.476 | ANT162 | -64.01 | -65.708 |
| ANT274 | -72.15 | -54.784 | ANT117 | -67.44 | -49.981 | ANT059 | -64.01 | -58.71 |
| ANT096 | -71.93 | -50.843 | ANT187 | -67.4 | -53.309 | ANT210 | -63.98 | -50.203 |
| ANT070 | -71.92 | -54.6 | ANT233 | -67.21 | -49.129 | ANT264 | -63.83 | -53.984 |
| ANT151 | -71.91 | -47.409 | ANT156 | -67.18 | -68.888 | ANT229 | -63.8 | -46.262 |
| ANT040 | -71.74 | -58.59 | ANT261 | -66.97 | -56.735 | ANT285 | -63.78 | -52.346 |
| ANT073 | -71.72 | -56.29 | ANT052 | -66.9 | -59.35 | ANT069 | -63.77 | -50.28 |
| **<u>ANT308</u>** | **<u>-71.56</u>** | **<u>-56.27</u>** | ANT011 | -66.77 | -50.199 | ANT181 | -63.71 | -53.046 |
| ANT157 | -71.52 | -61.406 | ANT176 | -66.76 | -53.694 | ANT197 | -63.6 | -69.074 |
| ANT225 | -71.44 | -50.818 | ANT099 | -66.74 | -51.45 | ANT168 | -63.56 | -52.247 |
| ANT138 | -71.21 | -52.437 | ANT275 | -66.69 | -47.826 | ANT041 | -63.52 | -46.674 |
| ANT133 | -71.19 | -60.864 | ANT263 | -66.61 | -51.594 | **<u>VIPhyb</u>** | **<u>-60.62</u>** | **<u>-51.007</u>** |
| ANT158 | -71.15 | -52.222 | ANT247 | -66.48 | -53.656 | ANT010 | -60.17 | -53.978 |
| ANT141 | -70.88 | -68.681 | ANT018 | -66.07 | -49.582 | ANT202 | -56.35 | -67.02 |
| ANT093 | -70.85 | -56.96 | **<u>VIP</u>** | **<u>-65.8</u>** | **<u>-52.61</u>** | ANT265 | -53.29 | -55.535 |
| ANT195 | -70.44 | -71.44 | ANT242 | -65.78 | -55.026 | ANT258 | -52.13 | -47.693 |
| ANT120 | -70.13 | -52.471 | ANT154 | -65.44 | -68.544 | ANT287 | -51.53 | 51.309 |
| ANT039 | -69.74 | -55.972 | ANT279 | -65.16 | -50.236 | ANT007 | -51.01 | -41.634 |
| ANT219 | -69.37 | -69.116 | ANT193 | -64.94 | -59.143 | ANT050 | -50.66 | -57.633 |
| ANT086 | -69.31 | -61.24 | ANT170 | -64.85 | -50.657 | ANT124 | -50.48 | -53.955 |
| ANT060 | -68.94 | -49.34 | ANT267 | -64.79 | -47.124 | ANT076 | -48.62 | -45.866 |
| ANT283 | -68.7 | -53.076 | ANT240 | -64.78 | -55.867 | ANT293 | -43.69 | -37.423 |
| ANT057 | -68.63 | -58.803 | ANT292 | -64.72 | -49.766 | ANT295 | -42.84 | -37.102 |
| ANT054 | -68.63 | -55.262 | ANT139 | -64.72 | -48.918 |  |  |  |

### Characterizing VIP Expression in C1498 Leukemia and P815 Mastocytoma Tumor Cells

Transcriptome analyses have indicated that VIP is overexpressed in a subset of AML patients (20, 21). We characterized VIP expression in murine C1498 myeloid leukemia and P815 mastocytoma cells using flow cytometry (Fig. 2A) and immunofluorescence (Fig. 2B, C). C1498 cells exhibited high VIP expression, hereon termed as VIP+ C1498 cells, whereas P815 cells displayed very low VIP levels. Additionally, P815 cells demonstrated low expression of the CD11b myeloid marker (Fig. 2B), which is consistent with previous reports (37).

**Figure 2.**
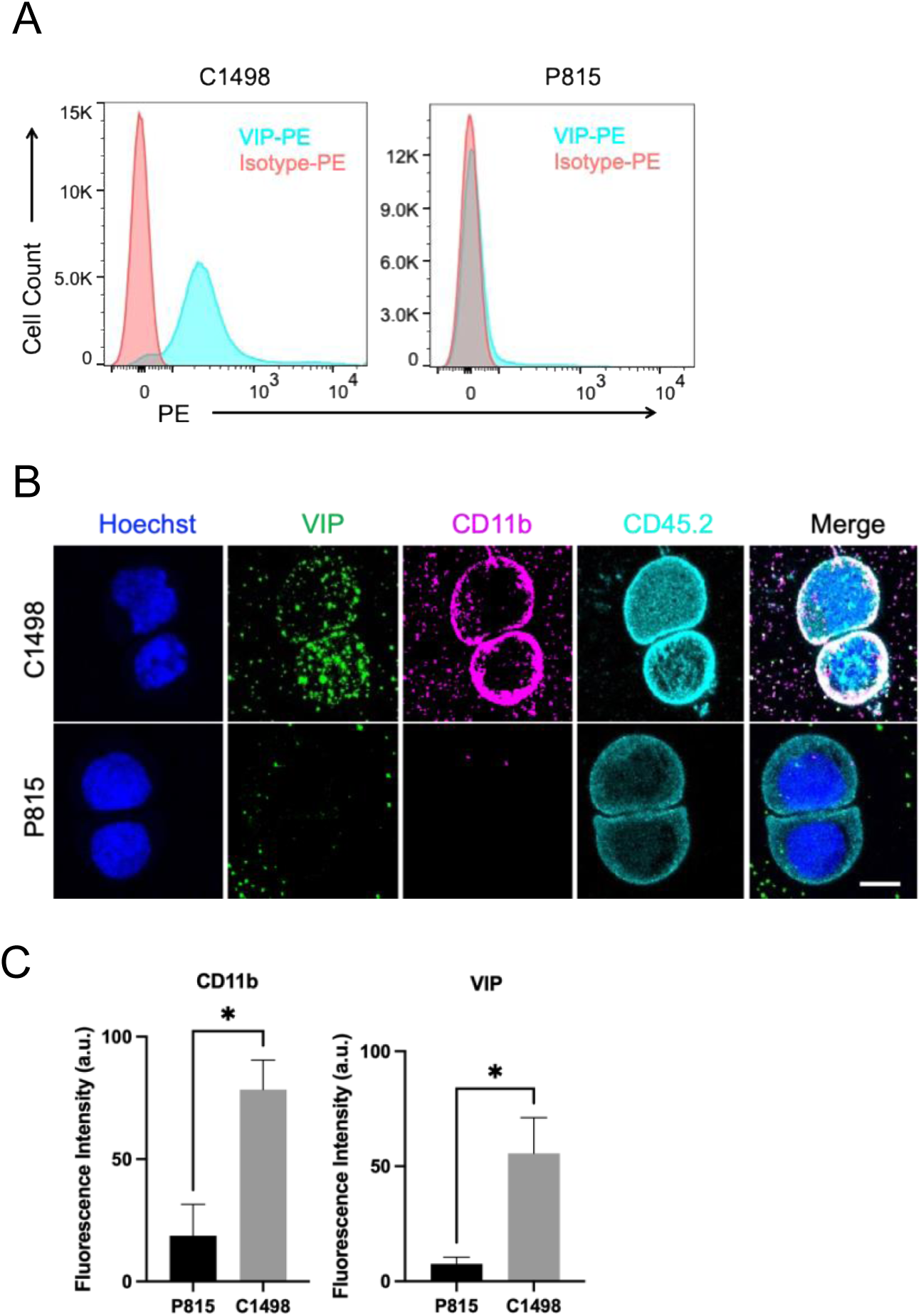
VIP expression in C1498 leukemia cells & P815 mastocytoma tumor cells. *(A)* Murine tumor cell lines P815 and C1498 cells were stained with VIP-PE or PE-isotype antibodies and then measured by flow cytometry. *(B)* Cytospins of C1498 and P815 cells were stained with fluorescently labeled antibodies to VIP (green), CD11b *(magenta)*, CD45.2 *(cyan),* and representative confocal *(x-y)* images are shown. Bar, 5μm. *(C)* Quantification of staining with antibodies to VIP and CD11b, unpaired t-test, * p<0.01.

### Novel VIP-ANT antagonists led to better survival in a murine model of acute myeloid leukemia

Given that autocrine VIP signaling by leukemia cells may exert direct growth-promoting autocrine effects on tumor cells (16, 22) and indirect paracrine effects on anti-cancer immunity by inhibiting T cells, we first tested the effect of ANT308 in a murine model of a VIP-secreting AML cell line, C1498. We injected CD45.2+ AML cells i.v. into congenic CD45.1+ B6 mice. Six days later, when mice had 2-6% leukemic blasts in their blood, they were treated with daily subcutaneous injections of 3 nM (10 μg) of VIP-ANT for 10 days. VIPhyb treatment led to ∼5% survival, whereas ANT308 & ANT195 treatment resulted in 40% survival (Fig. 3A-C). Negative control mice treated with daily injections of PBS or VIP Fully SCRAM (in which all VIP amino acids 1-28 are randomly scrambled) all died within 40 days after tumor inoculation (Fig. 3 A-C). VIP-knock-out (B6 background) mice had >60% survival (p<0.0001, compared to VIP Fully SCRAM), confirming the role of VIP-signaling in regulating anti-cancer immunity. Monitoring the content of CD45.2+ leukemia cells in blood samples obtained from day 0 to day +25 of treatment showed similar levels of leukemia cells on day +6 at the initiation of therapy (median 3.5%) (Fig. 3 D-F). Leukemia frequencies increased to >15% leukemia blood cells on day 20 in control mice treated with VIP Fully SCRAM. In contrast, treatment with ANT308 or ANT195 significantly reduced the median content of leukemia cells content in blood to 2.06% or 9.84%, respectively. Thus, ANT308 & ANT195 showed the most potent anti-leukemia activities in the VIP+ C1498 murine model of AML.

**Figure 3.**
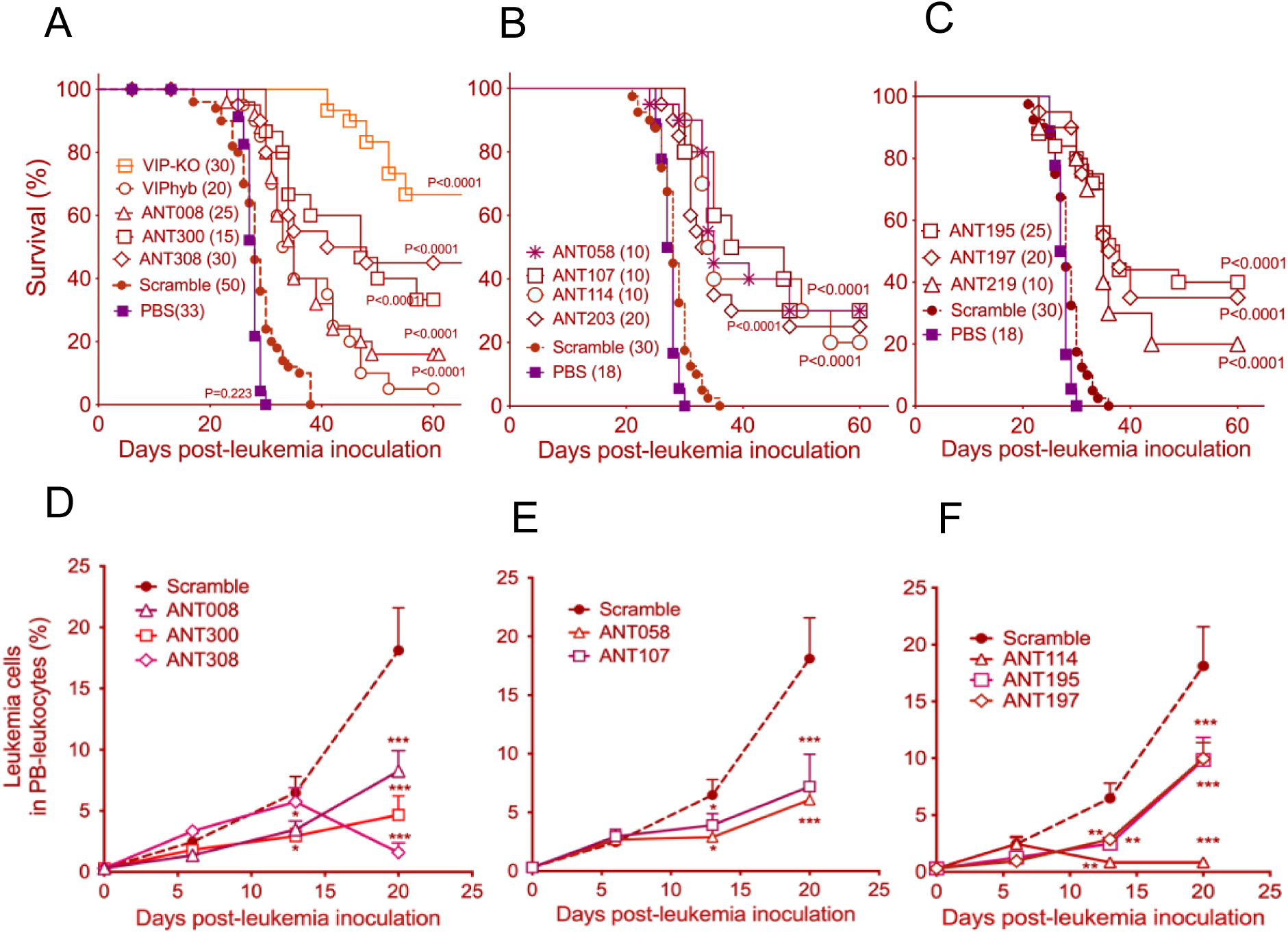
Treatment with novel VIP-ANT antagonists improved survival compared to VIPhyb in a murine model of acute myeloid leukemia. B6 mice were injected intravenously with 1 x 10^6^ C1498 cells iv on day 0, then treated with 10μg of various VIP-ANTs injected subcutaneously once daily from day +6 through day +12. *(A-C)* Survival curves of mice treated with various VIP-ANTs, (N=10-30). The mice were pooled from 3-4 replicated experiments and the survival was analyzed with log-rank test. VIP-ANTs were grouped based on their number of mutations: panel A are VIP-ANTs with 1-3 a.a. substitutions in addition to VIPhyb mutations; VIP-ANTs in B contains 3-6 substitutions in addition to VIPhyb mutations; VIP-ANTs in C contains 5-6 substitutions in addition to VIPhyb mutations. *(E-G)* Flow cytometry measurement of C1498 leukemia cells in peripheral blood from day 0 to day 25 (n=10). The data were pooled from two replicated experiments. Significance among the groups were analyzed with One-Way ANOVA, post-hoc multiple comparison with Tukey process by SPSS version 29. *p<0.05, **p<0.01 and ***p<0.001.

### VPAC1 and VPAC2 receptor docking scores significantly correlated with leukemic mice survival rates and T cell proliferation rates

Next, to test if VIP-ANTs with higher predicted receptor binding affinities would better activate T-cells or exhibit higher anti-leukemic efficacies, we measured the proliferation of luciferase+ B6 splenic T-cells following TCR-activation in the presence of 3 μM VIP-ANTs, then plotted normalized T-cell proliferation rates with the docking scores of VIP-ANTs to VPAC1, VPAC2 or the sum (Fig. 4 A-C). Regression analysis on scatter plots and R^2^ suggested that the proliferation of murine T-cells best correlated with the sum of the docking scores to VPAC1 and VPAC2 (R^2^=0.6368). Then, we performed a regression analysis comparing the VPAC1 and VPAC2 docking scores with the fraction of mice cured of C1498. We found that day 60 leukemia-free survival best correlated with the sum of the docking scores to VPAC1 and VPAC2 (R^2^=0.7282) (Fig. 4 D-F). These results indicate that VIP signaling through both VPAC1 and VPAC2 suppresses T cell activation and anti-leukemia activities.

**Figure 4.**
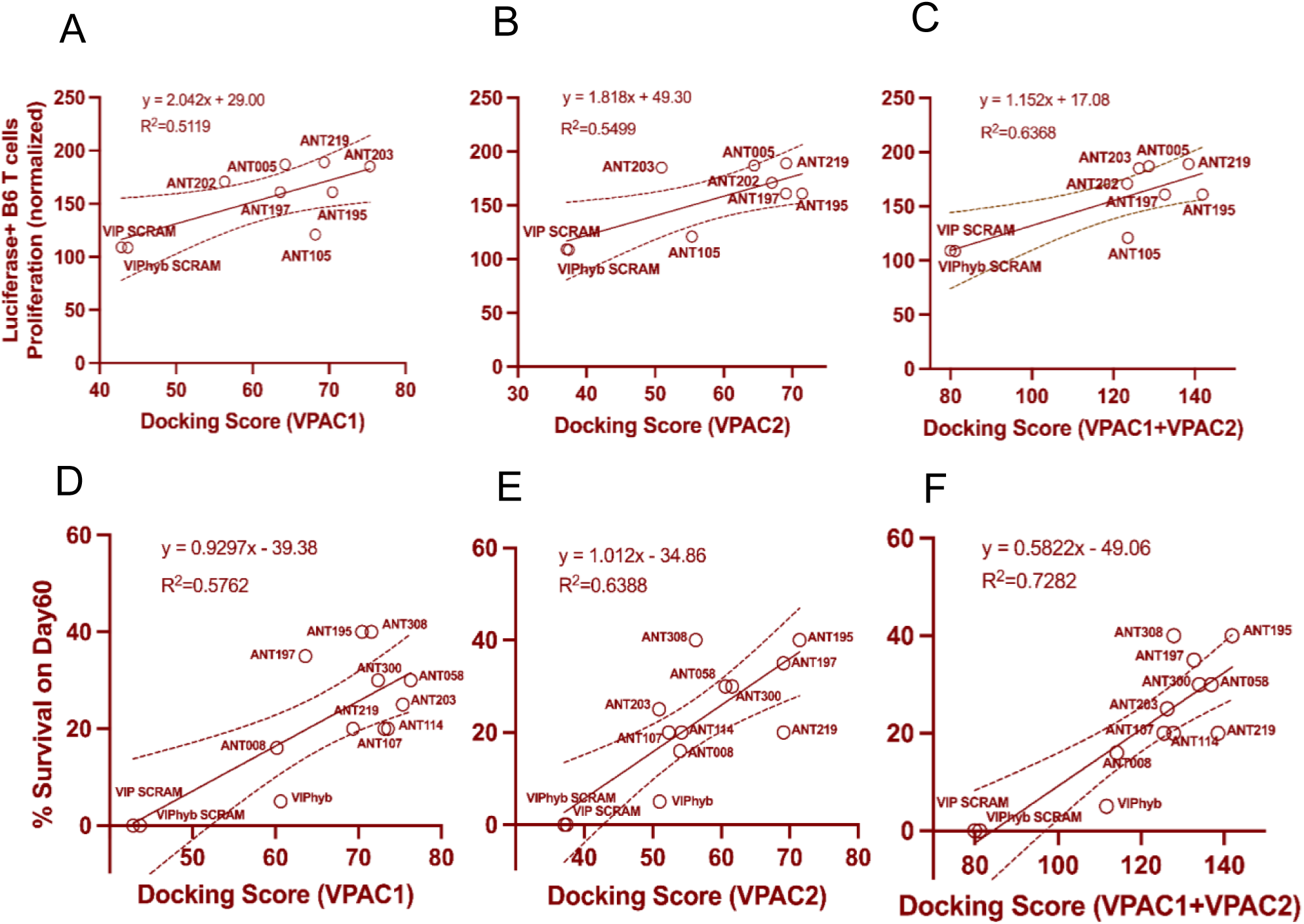
VPAC1 and VPAC2 receptor docking scores correlated with mouse T cell proliferation rates and survival of VIP-ANT-treated leukemic mice. *(A-C)* Luciferase B6 splenic T cell proliferation levels in culture wells stimulated with anti-CD3e and IL2 and treated with 3uM of various VIP-ANTs for 72 hours, normalized to proliferation in untreated wells, were plotted against VIP-ANTs docking scores (dependent on binding affinities of antagonists to VPAC1, VPAC2, or both receptors) predicted by *in silico* modeling. R-squared values show correlation levels between T-cell proliferation and docking score for the receptor types. *(D-F)* % Survival of B6 mice 60 days after inoculation with C1498 tumor cells was recorded for mice treated with various VIP-ANTs. R-squared values show the correlation levels between the survival of mice inoculated with tumor cells and VIP-ANTs docking scores to VPAC1, VPAC2, or the sum of both scores.

### ANT308 enhances the activation and proliferation of human T cells and decreases CREB phosphorylation

We tested whether the two VIP-ANTs, ANT308 & ANT195, which resulted in the highest anti-leukemic activity *in vivo* (Fig. 3), could potentiate human T cell activation. To directly measure the effects of autocrine signaling from VIP secreted by activated T cells, we purified T cells from different healthy volunteer donors, incubated them with plate-bound anti-CD3 antibody, and measured their activation in the presence of VIP-ANTs or VIP Fully SCRAM (Fig. 5A-C). 3 μM ANT308 significantly boosted CD69 expression on both CD4+ and CD8+ human T cells without affecting T cell viability. In contrast, ANT195 potently activated T cells at 1 μM but was toxic to T cells at 3 μM, with only ∼30% viability after 24hr culture. Hence, we chose ANT308 as the most promising VIP-receptor antagonist for further studies. Recognizing the significant inter-subject variability in the level of T cell activation *in vitro*, we used pooled human T cells from 4-5 different donors to determine the in vitro EC_50_ of ANT308 based on measuring levels of CD69 expression on CD4+ or CD8+ T cells treated with increasing concentrations of ANT308 (Fig. 5 D-E). VIPhyb causes a half-maximal inhibition of VIP binding to VIP receptors on lymphocytes at 5 μM (28). ANT308 significantly increased T cell activation with an EC_50_ around 0.3-0.4 μM, in agreement with the higher predicted docking score. Our previous work showed that VIPhyb enhances T cell proliferation and reduces CREB phosphorylation in CD4+ and CD8+ T cells following VIP stimulation (23). In line with the mechanism of action of VIP-ANT, 3 or 10 μM ANT308 significantly increased Ki67 expression on the CD3+ T cell population (Fig. 5F) and 10 μM ANT308 decreased CREB phosphorylation levels in purified human T cells upon stimulation with 1 or 10 nM VIP (Fig. 5G).

**Figure 5.**
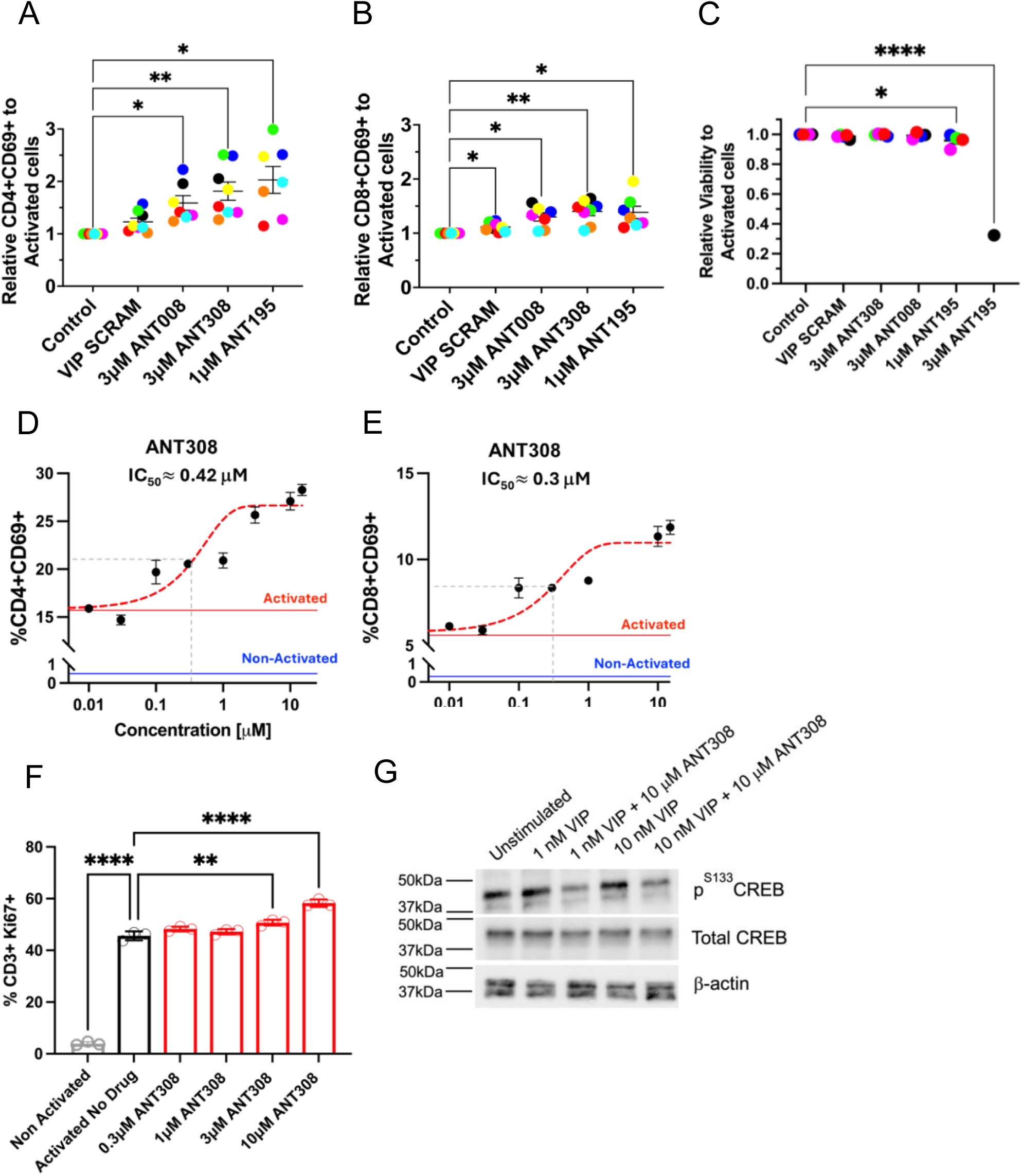
ANT308 enhanced activation, proliferation, and decreased CREB phosphorylation in human T cells from healthy donors. ANT008 (3 μM), ANT308 (3 μM), VIP SCRAM (3 μM) or ANT195 (1,3 μM) were incubated with isolated T cells from different donors (each symbol color denotes individual healthy donors) in the presence of plate-bound anti-CD3 antibody & human IL-2 (30 IU/ml). 24 hrs following incubation, cell subsets were examined for (*A,B)* CD69 expression levels and (*C)* cell viability, one-way ANOVA, *p<0.05, **p<0.01, ****p<0.0001. (*D,E)* Isolated & pooled T cells from 3-5 healthy donors were incubated with increasing concentrations of ANT308, in the presence of soluble human CD3/CD28/CD2 T cell activator & human IL-2 (50 IU/ml) for 48hrs. %CD4+CD69+, %CD8+CD69+ were then quantified by flow cytometry and concentration-dependent curves were generated and used to determine EC_50_ values. (*F)* %CD3+Ki67+ population in response to increasing concentrations of ANT308. The data represent the means ± SD of three independent experiments in triplicate. one-way ANOVA. **p<0.01, ****p<0.0001 (*G)* Western blot showing CREB phosphorylation in isolated pooled human T cells treated with 1 or 10nM VIP in the presence or absence of 10 μM ANT308.

### ANT308 significantly increases GZMB, and Perforin expression in CD8+ AML patients’ T-cells

Perforin and granzyme B expression were measured in a pool of CD8+ T cells isolated from 4-5 AML patients’ peripheral blood mononuclear cells (PBMC). A dose-dependent increase in perforin expression was seen in CD8+ T cells from AML patients with increasing concentrations of ANT308 (Fig. 6 A & B). Intriguingly, perforin expression remained unchanged in T cells from healthy donors after blocking VIP signaling (data not shown). A moderate increase in the level of granzyme B expression in CD8+ AML T cells was detected following 3 μM ANT308 treatment (Fig. 6 C & D). The results indicate that blocking VIP-receptor signaling specifically triggers the perforin-granzyme pathway in T cells from AML patients, supporting the potential of VIP-ANT to augment T cell response for tumor surveillance in human AML cancer.

**Figure 6.**
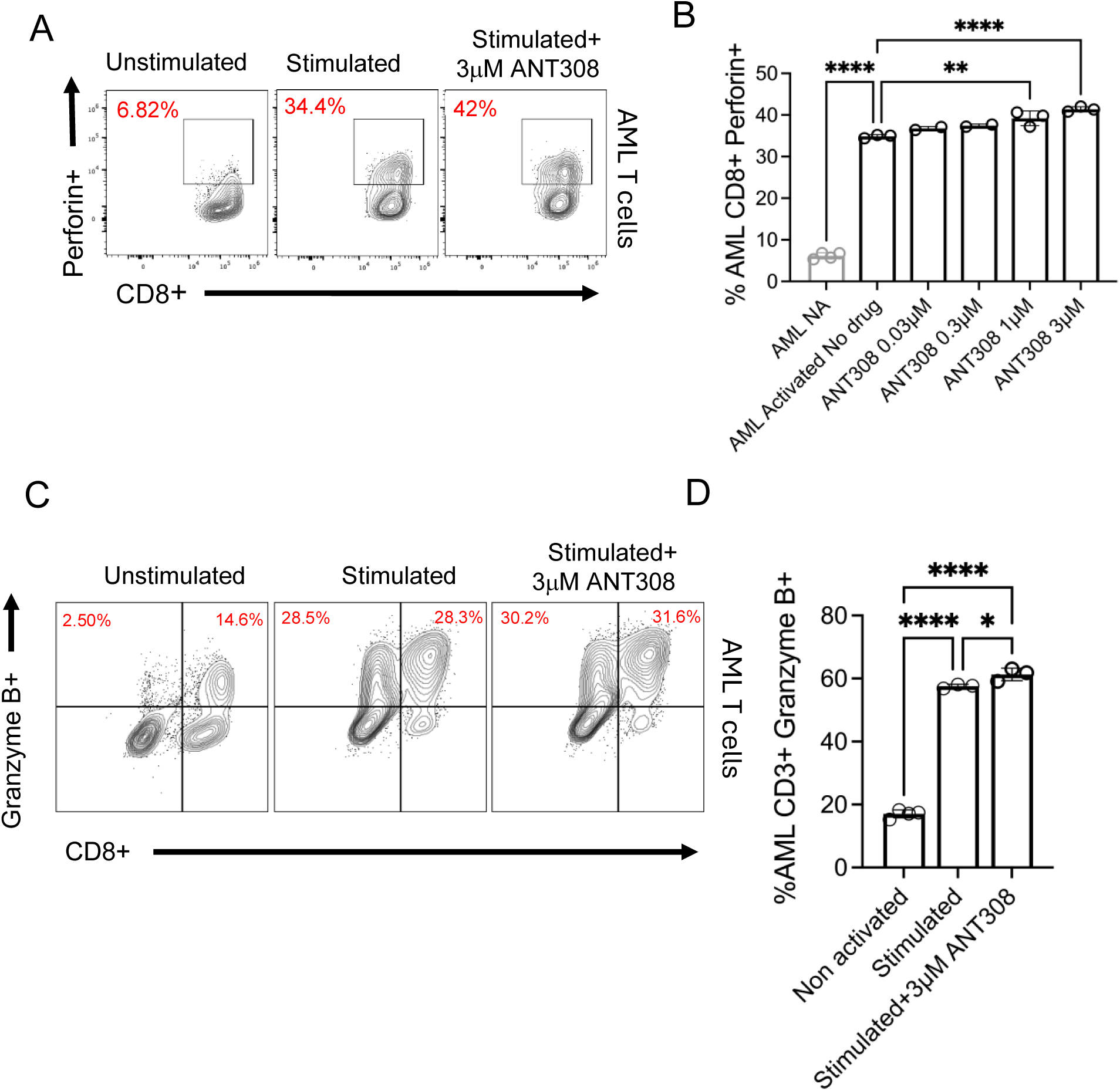
ANT308 increased Perforin and Granzyme B expression in T cells from AML patients. Four AML patients’ T cells were isolated and pooled, treated with various concentrations of ANT308 for 48 hours in the presence of αCD3/CD28 activator & IL-2. (*A-B)* %CD8+Perforin+ T cells were quantified by flow cytometry, and representative flow cytometry plots of cells were shown, n=3. (*C-D)* %CD3+ Granzyme B+ T cells were quantified by flow cytometry (n=3), ANOVA, *p<0.05, ****p<0.0001

### The anti-leukemia effect of ANT308 is dose and schedule dependent

Next, we sought to determine the efficacy of ANT308 *in vivo* with various dosages and administration schedules. B6 mice were inoculated with C1498 and treated with either 30μg VIP Fully SCRAM or 10, 30, or 100μg of ANT308 starting seven days after tumor inoculation, daily for 2 weeks. Higer dosages correlate with better survival (Fig. 7A). In a separate experiment, B6 albino mice were inoculated with C1498ff cells and treated with a high dose of ANT308 (100 μg, equal to 30 nM) twice daily, once daily, or every other day, starting seven days post tumor inoculation and continuing for 2 weeks. We found that twice daily treatment yielded the highest survival (65%), followed by once daily and every other day treatment (Fig. 7B). Serial bioluminescent imaging (BLI) showed the largest reduction in C1498ff leukemia burden with ANT308 administered twice daily (Fig. 1D). We found that cumulative dosages positively correlated with survival (Fig. 7C, results summarized from Fig. 7A & 7B). Thus, our data confirmed a dose and schedule dependency of ANT308 treatment. Notably, high-dose ANT308 did not produce any observable toxic effects in the weight or behavior of treated mice, consistent with our previous report documenting the absence of organ pathology or abnormal blood tests in VIP ANT-treated mice(5).

**Figure 7.**
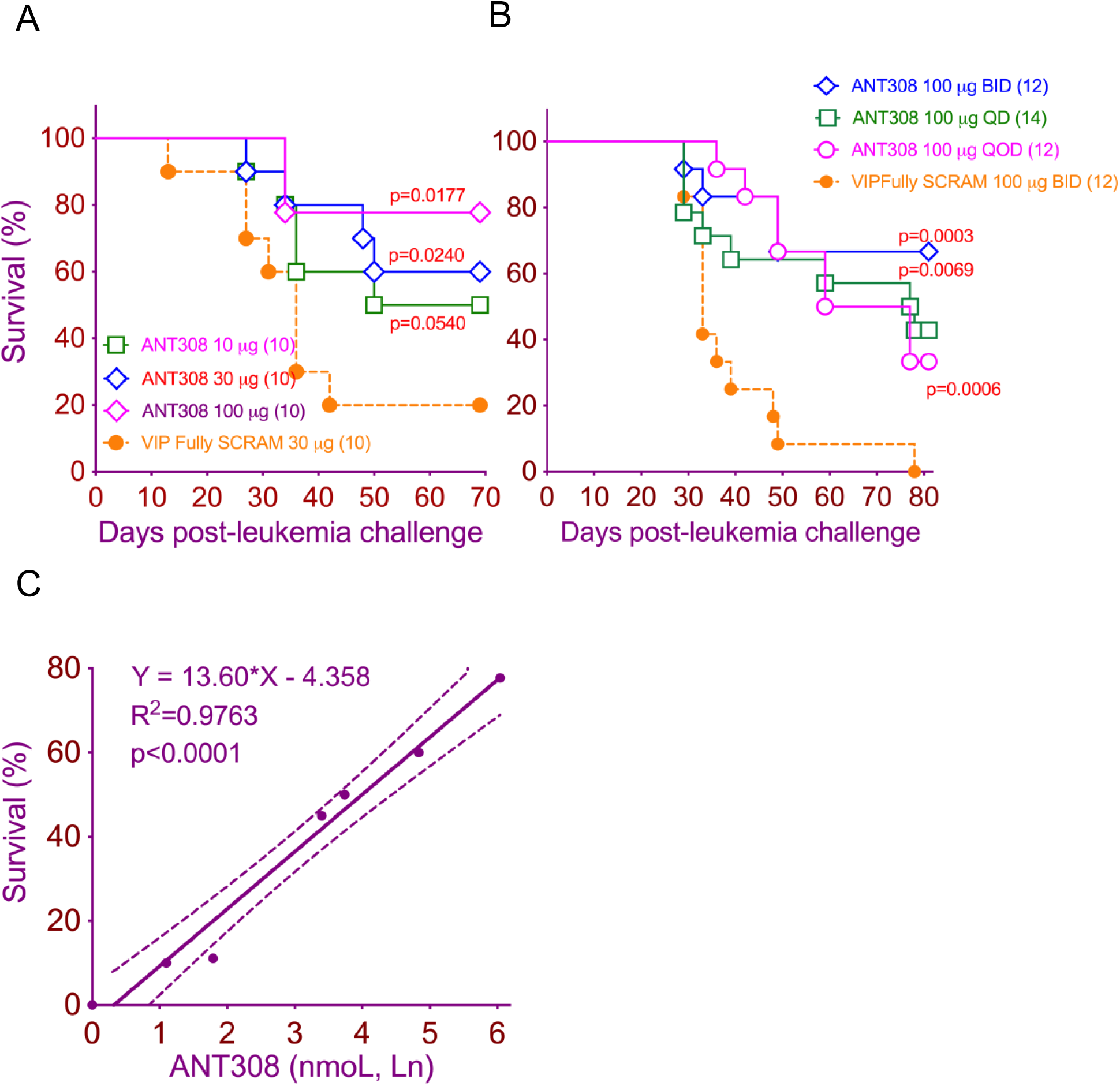
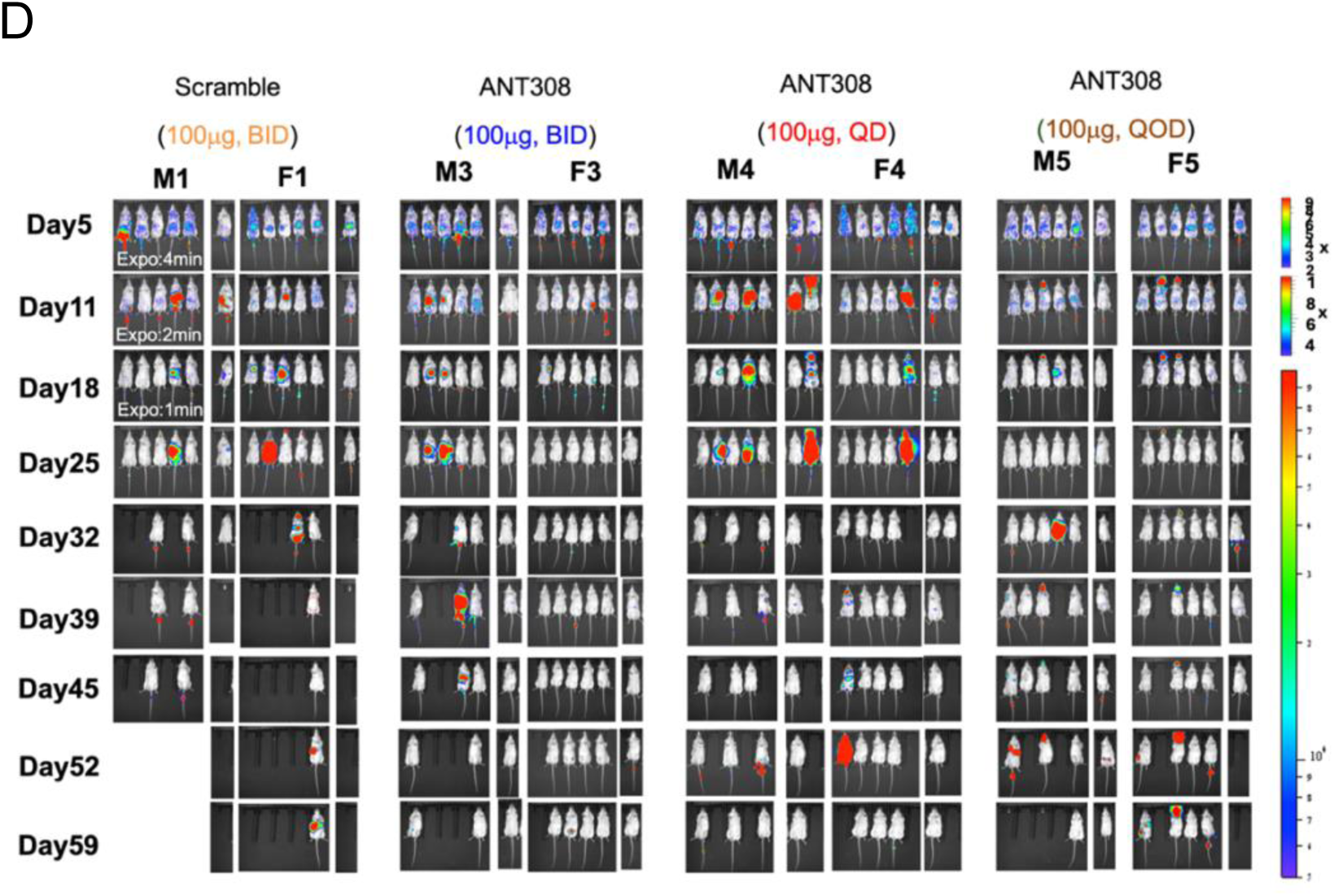
The anti-leukemia effect of ANT308 is dose and schedule dependent. *(A)* Survival curves for B6 mice injected with 1 x10^6^ C1498 on day 0, and treated with either 10, 30, 100 μg of ANT308, or 30 μg VIP Fully SCRAM starting 7 days after tumor inoculation and continuing daily for 2 weeks. *(B)* Survival curves for B6 albino mice (male and female) inoculated with 1 x10^6^ C1498ff cells, treated with 100 μg ANT308 on different schedules: BID (twice daily), QD (daily), or QOD (every other day) starting 7 days after tumor inoculation and continuing daily for 2 weeks. Control mice received twice-daily injections of 100 μg VIP Fully SCRAM. Pairwise survival differences between ANT308-treated and control groups were evaluated using the log-rank test. *(C)* Survival plotted against cumulative ANT308 dosages (natural logarithm), analyzed by simple linear regression. *(D)* Tumor progression of male (M1,3,4,5) and female mice (F1,3,4,5) from panel B, monitored by BLI over 60 days.

### Novel VIP-ANT antagonists induced protective anti-leukemia immune memory among mice with VIP-secreting AML and VIP-non-secreting myeloid malignancies

Our previous study showed VIP receptor antagonists conferred protective immunity to tumor re-challenge (5, 38). To test if ANT308 treatment yields long-term anti-leukemia protection, we first tested whether VIP-receptor antagonists would be effective in a VIP-non-secreting myeloid cancer cell line, P815 mastocytoma (Fig. 8A, 8B). Our *in vitro* screen demonstrated T-cell intrinsic effects of blocking VIP-receptor signaling on cultures of purified T cells (Fig 5). We hypothesized that blocking VIP signaling in VIP-non-secreting myeloid cancers could enhance T-cell-mediated autologous anti-leukemia responses in mice with a VIP-non-secreting cell line by blocking autocrine VIP signaling among T cells or paracrine effects of VIP synthesized by other non-malignant immune cells in the tumor microenvironment. We established P815 mastocytomas in DBA/2 mice by s.c. injection of 1 x 10^5^ tumor cells. Six days later, treatment with 10 daily s.c. injections of ANT308 on the opposite flank commenced. ANT308 treatment led to complete regression of established tumors in 50% of ANT308-treated mice compared with 100% tumor progression and death among mice treated with injections of a control, scrambled-sequence peptide (Fig. 8A). Thus, ANT308 induced potent anti-leukemia responses in both VIP-secreting (C1498, Fig. 3, Fig. 7) and VIP-non-secreting myeloid tumors (P815, Fig. 8A&B). Next, we tested whether mice rendered tumor-free after P815 or C1498 injections following VIP-ANT treatment had developed protective immunological memory. B6 mice that survived initial inoculation with P815 (Fig. 8A) or C1498 inoculation (1 x 10^6^ leukemia cells; Fig. 8C; select groups from Fig. 3) were re-challenged with a 2-fold higher dose of P815 (2 x 10^5^) or C1498 (2 x 10^6^), respectively. Prior VIP-ANT treatment led to >60% survival in the P815 rechallenge model (Fig. 8B) and >80% survival in the C1498 rechallenge model (Fig. 8D), consistent with our previous report that VIP signaling blockade activates and expands memory CD8+ T cells in a murine AML model (38).

**Figure 8.**
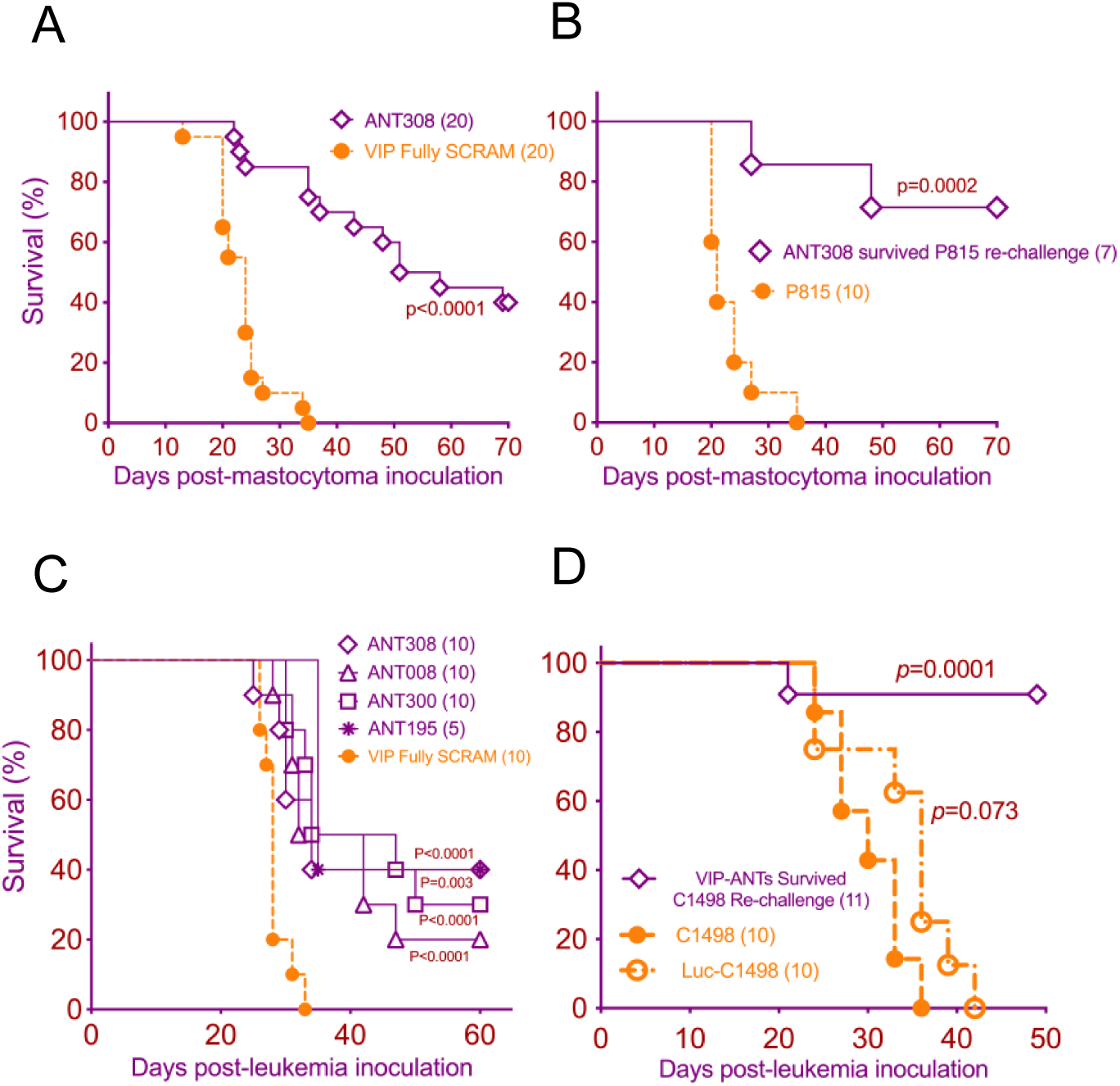
Novel VIP-ANT antagonists induced protective immunological memory against P815 and C1498. (*A)* Survival of DBA/2 mice following s.c. injection of 1 x 10^5^ P815 mastocytoma cells and treatment with once daily injections of 100 μg ANT308 or VIP fully scrambled peptide on days +6 to +16. (*B)* Surviving mice from Fig. 8(A) and naïve DBA/2 mice received a s.c. injection of 2 x 10^5^ P815 cells (on the opposite flank for re-challenged mice). *(C)* Depicts the survival of B6 mice with C1498 leukemia after treatment with ANT308, ANT008, ANT300, or ANT195 that were used for the tumor re-challenge experiment (from Fig. 3). *(D)* Tumor-free mice from Fig. 8C were combined and re-challenged with 2 x 10^6^ C1498 cells vs. naive B6 mice that received either C1498 or luciferase-expressing C1498-ff tumor cells. The significance of survival differences between the re-challenge and C1498-ff groups with the C1498 group are measured with the log-rank test.

### VIP, ANT308, VIPhyb binding to VPAC receptors predicted by AlphaFold

To gain insight into the molecular interactions of VIP agonists and antagonists with VIP receptors, we used AlphaFold for conformational analyses of agonist- and antagonist-bound complexes between VIP, VIPhyb and ANT308 with VIP receptors VPAC1 and VPAC2 (39–42). As shown in Fig. 9A/D & Fig. 9G/J, the N-terminus of VIP is deeply inserted into the transmembrane bundle of alpha helices of VPAC1 and VPAC2 receptors, with H-bonds formed between VIP T11 & VPAC1 D287, and VIP T7 & VPAC1 K2.67; and a salt bridge between VIP D3 & VPAC1 R2.60, consistent with a previous cryo-EM study(43). VIP and PACAP27 N-termini conserved residues form identical interactions with VPAC2: Hydrophobic contacts between VIP H1 & VPAC2 I213^3.40b^, F6 & Y123^1.36b^, Y130^1.43b^, H-bonds between VIP S2 & VPAC2 E360^7.42b^, VIP T7 & K179^2.67b^, D8 & N275^ECL2^, salt bridge between D3 & VPAC2 R172^2.60b^, R12 & D276^ECL2^(Fig. 9G &J; (44)). In contrast, the substitution of the neurotensin sequence in the N-terminus led to a conformational bend in ANT308 & VIPhyb (Fig. 9B&H; Fig. 9C&I). Similar interactions between VIP-ANT and VIP-R were noted in VIPhyb & ANT308 bound to murine VPAC1 and VPAC2 as predicted by AlphaFold (Fig.S3). As seen with the human VIP-R receptors, the N-terminus of the VIP-ANT are orthogonal to the axis of the alpha-helical C terminal sequences. Differences in the binding interactions between the N terminus of VIP versus the VIP-ANT might alter allosteric conformational changes in the receptors that regulate intracellular heterotrimeric GTP-binding protein (G protein) activation (45). However, the AlphaFold modeling does not show significant conformational changes in the intracellular region of human VPAC1 comparing binding of VIP versus ANT308 (Fig.S1), which might be due to a low Predicted Local Distance Difference Test (pLDDT) of less than 50 in predicting the intracellular domains of VPAC1/2 with or without bound peptide ligands.

**Figure 9.**
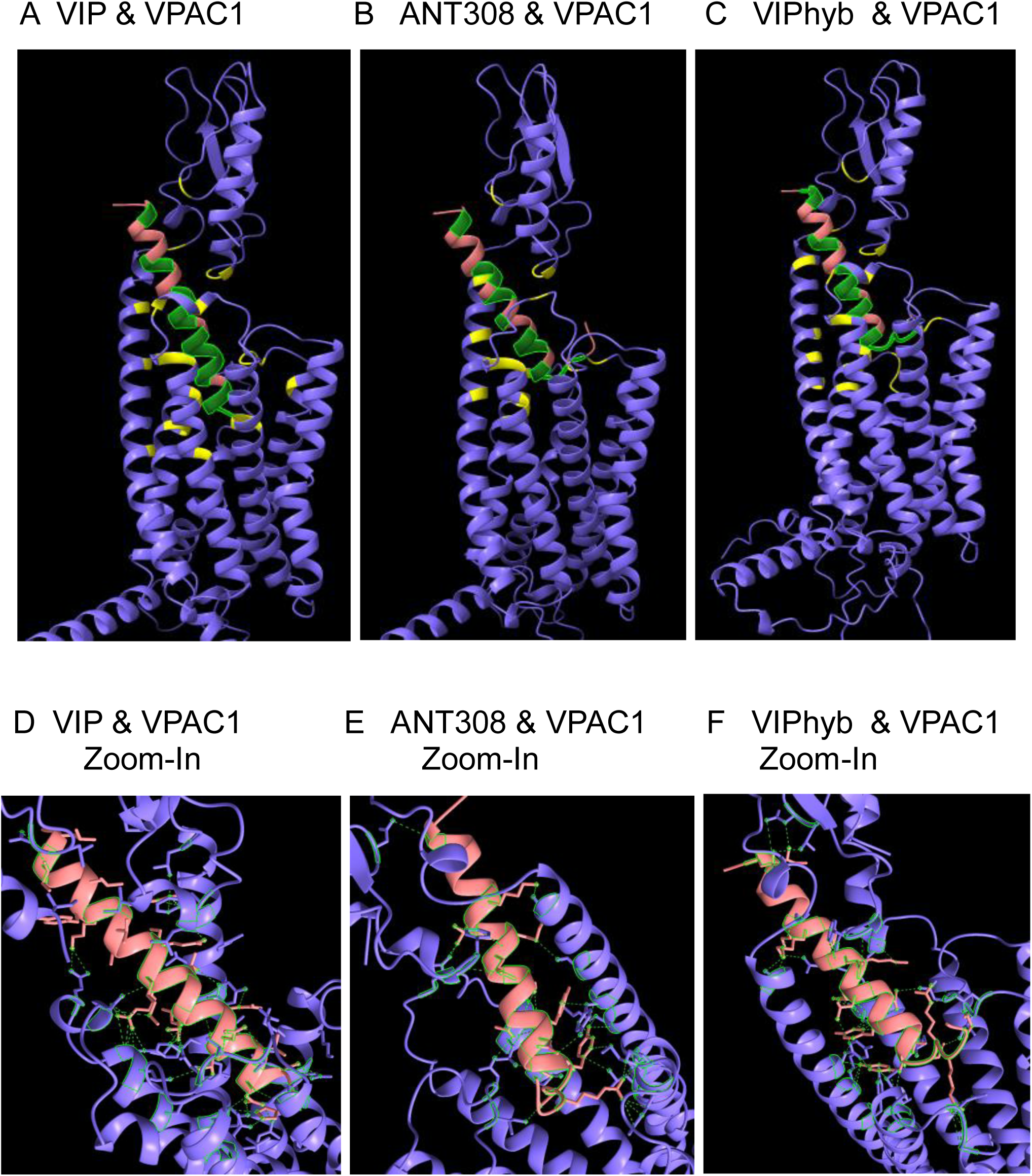
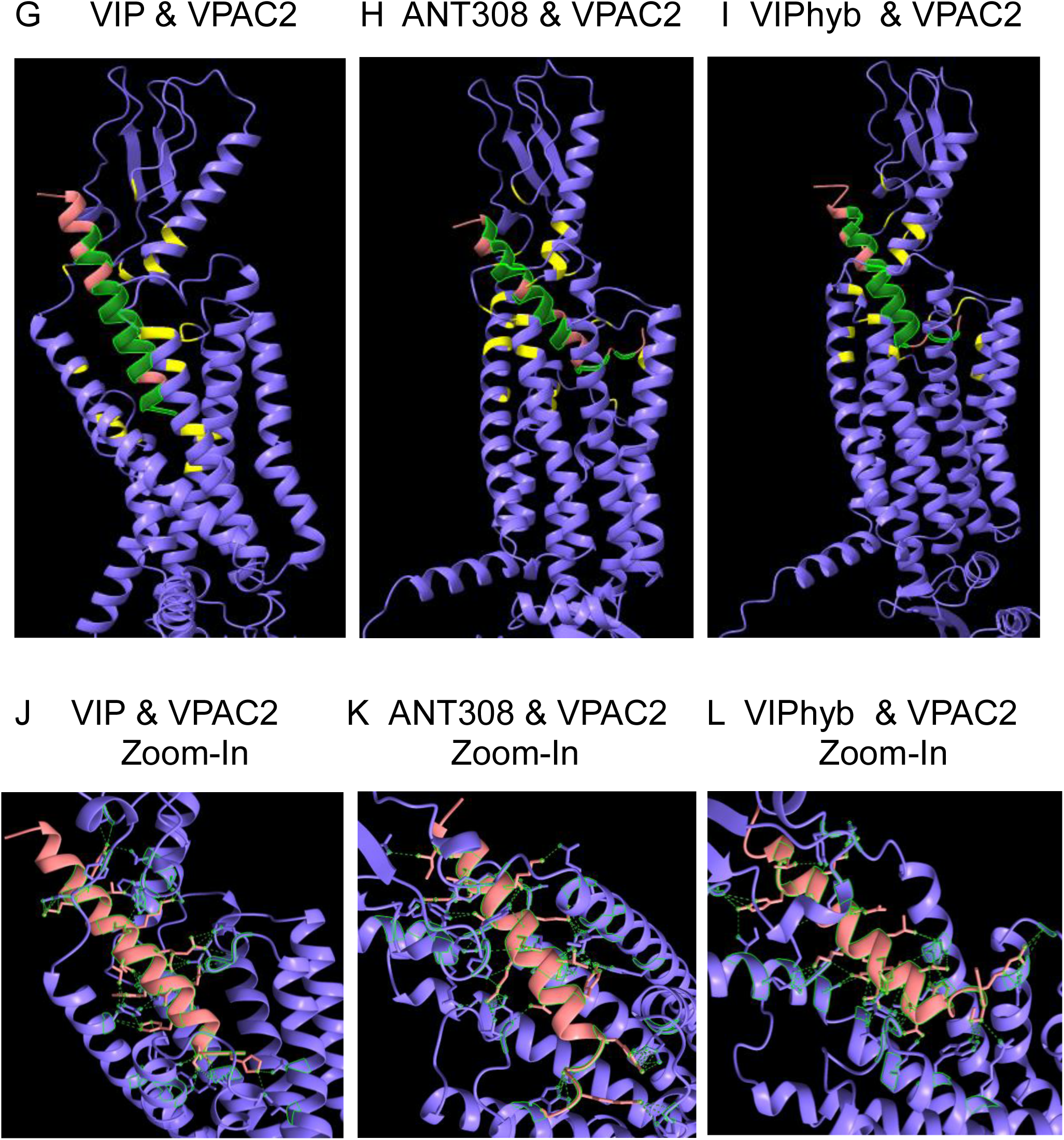
The binding of peptide antagonists to human VPAC1, VPAC2 as represented by AlphaFold. Secondary structure comparison of human VPAC1 binding to VIP, ANT308, and VIPhyb. Human VPAC1 (purple multi-domain complex, VIP binding interfaces highlighted in yellow). *(A)* VIP, *(B)* ANT308, *(C)* VIPhyb (pink peptide helices and beta sheets, VPAC1 binding interfaces highlighted in green). *(D-F)* Zoom-In view of VIP, ANT308 or VIPhyb binding to the VPAC1 receptor, pseudo bonds are highlighted in green. *(G-L)* Human VIP-VPAC2, ANT308-VPAC2, VIPhyb-VPAC2 complex solved by AlphaFold. Secondary structure comparison of human VPAC2 binding to VIP, ANT308, and VIPhyb. Human VPAC2 (purple multi-domain complex, VIP binding interfaces highlighted in yellow). *(G)* VIP, *(H)* ANT308, *(I)* VIPhyb (pink peptide helices and beta sheets, VPAC1 binding interfaces highlighted in green). *(J,K,L)* Zoom-In view of VIP, ANT308 or VIPhyb binding to the VPAC2 receptor, pseudo bonds are highlighted in green.

The amino acid side chains of the C-terminus of the VIP-ANT form extensive network of engagement with residues in the receptors, and the effect of sequential amino acid substitutions in ANT008 and ATN300 to create ANT308 (Table 3 & 4) (Fig. S2). Notably, some of the substituted residues form new interactions with residues in the receptors not seen in VIPhyb bound to VAPC1, i.e., N9D in ANT300 and ANT308 interact with Y98 in VPAC1. In contrast, other substituted residues led to subtle conformation changes in the peptide, with the single substitution in ANT008 leading to an adjacent residue interacting with a new residue in the VPAC2 receptor (K15 interacting with F59), and the two substitutions in ANT300 and ANT308 (D8S and N9D) led to a new predicted interaction of M17 interacting with T95 in VPAC1. D8S, N9D, S25L in ANT308 also cause different engagement with mouse VIPR1/2 (Supplement Table 1, 2)

**Table 3.** Amino acid residues involved in Human VPAC1 and peptide interactions predicted by ChimeraX. Differences from VIP sequence are indicated by *<u>bold,</u> <u>underlined, and italicized</u>* letters. Residue numbers were counted directly within the Human VPAC1 (PDB ID:8e3z) sequence (Protein Data Bank, Europe).

| VIPhyb | Human VPAC1 | ANT008 | Human VPAC1 | ANT300 | Human VPAC1 | ANT308 | Human VPAC1 |
| --- | --- | --- | --- | --- | --- | --- | --- |
| <u>**K1**</u> |  | <u>**K1**</u> |  | <u>**K1**</u> |  | <u>**K1**</u> |  |
| <u>**P2**</u> | N249 | <u>**P2**</u> |  | <u>**P2**</u> |  | <u>**P2**</u> |  |
| <u>**R3**</u> | P320, D321, R319 | <u>**R3**</u> | D321,N249 | <u>**R3**</u> | D321, P325, F323, <u>**K328**</u> | <u>**R3**</u> | P325, K328 |
| <u>**R4**</u> | N249 | <u>**R4**</u> |  | <u>**R4**</u> | N249 | <u>**R4**</u> |  |
| <u>**P5**</u> | I248 | <u>**P5**</u> |  | <u>**P5**</u> |  | <u>**P5**</u> | I248 |
| <u>**Y6**</u> | Y98, K102, M32, L333, Y105 | <u>**Y6**</u> | Y98, M329 | <u>**Y6**</u> | Tyr98, K102, M329 | <u>**Y6**</u> | Y98, K102, L333, M329 |
| T7 | K154 | T7 |  | T7 | K154 | T7 | K154, L158 |
| D8 |  | D8 |  | <u>**S8**</u> | N246, T247 | <u>**S8**</u> |  |
| N9 |  | N9 |  | <u>**D9**</u> | Y98 | <u>**D9**</u> | Y98 |
| Y10 | F159, Y98 | Y10 | F159, Y98 | Y10 | Y98, F159 | Y10 | Y98 |
| T11 | D246, F159 | T11 |  | T11 | D246 | T11 | D246, F159 |
| R12 |  | R12 |  | R12 |  | R12 |  |
| L13 | Y98 | L13 | Y98, T95 | L13 |  | L13 |  |
| R14 | F159, Q166 | R14 | F159 | R14 | F159, Q166 | R14 | F159 |
| K15 | Q166 | K15 | D165, S164 | K15 |  | K15 |  |
| Q16 | L51, F52 | Q16 | L51, D91, F52 | Q16 | L51, F52 | Q16 | L51, F52 |
| M17 |  | M17 |  | M17 | T95 | M17 | T95 |
| A18 |  | A18 |  | A18 |  | A18 |  |
| V19 |  | V19 | I48 | V19 | I48 | V19 | Q166, F52 |
| K20 | A88, L83 | K20 | A88 | K20 |  | K20 | A88 |
| K21 | E163 | K21 |  | K21 |  | K21 |  |
| Y22 |  | Y22 |  | Y22 |  | Y22 |  |
| L23 |  | L23 |  | L23 |  | L23 |  |
| N24 |  | N24 |  | N24 |  | N24 |  |
| S25 |  | <u>**L25**</u> |  | S25 |  | <u>**L25**</u> |  |
| I26 | N28 | I26 | N28 | I26 | N28 | I26 | N28 |
| L27 | L29 | L27 |  | L27 |  | L27 |  |
| N28 |  | N28 |  | N28 |  | N28 |  |

**Table 4.** Amino acid residues involved in Human VPAC2 and peptide interactions predicted by ChimeraX. Differences from VIP sequences are indicated by *<u>bold,</u> <u>underlined, and italicized</u>* letters. Residue numbers were counted directly within the Human VPAC2 (PDB ID:7vqx) sequence (Protein Data Bank, Europe).

| VIPhyb | Human VPAC2 | ANT008 | Human VPAC2 | ANT300 | Human VPAC2 | ANT308 | Human VPAC2 |
| --- | --- | --- | --- | --- | --- | --- | --- |
| <u>**K1**</u> |  | <u>**K1**</u> |  | <u>**K1**</u> |  | <u>**K1**</u> |  |
| <u>**P2**</u> | W258, R262 | <u>**P2**</u> | W258 | <u>**P2**</u> |  | <u>**P2**</u> |  |
| <u>**R3**</u> | I328, Q333 | <u>**R3**</u> | I328, Q333 | <u>**R3**</u> | W258, P325 | <u>**R3**</u> | P325 |
| <u>**R4**</u> |  | <u>**R4**</u> | D253 | <u>**R4**</u> | D253 | <u>**R4**</u> |  |
| <u>**P5**</u> |  | <u>**P5**</u> |  | <u>**P5**</u> | W258 | <u>**P5**</u> | W258 |
| <u>**Y6**</u> | Y100, K104 | <u>**Y6**</u> | Y100, K104, L338 | <u>**Y6**</u> | Q333, E337 | <u>**Y6**</u> | Q333, E337 |
| T7 | Y161, K156 | T7 | Y161, K156 | T7 |  | T7 |  |
| D8 | N252, D250 | D8 | N252, D250 | <u>**S8**</u> |  | <u>**S8**</u> |  |
| N9 | Y100 | N9 | Y100 | <u>**D9**</u> |  | <u>**D9**</u> |  |
| Y10 | Y161, Y100 | Y10 | Y100, Y161 | Y10 | Y107, Y100 | Y10 | Y107, Y100 |
| T11 | D250, Y161 | T11 | D250, Y16, R3 | T11 | D250 | T11 | D250 |
| R12 | D253 | R12 | D253 | R12 |  | R12 |  |
| L13 | Y100 | L13 | I97 | L13 |  | L13 | K96 |
| R14 | S164, Y16, E1 | R14 | E1, Y161 | R14 | D157, Y100, Y161, Y107, K104 | R14 | Y161, D157, Y100 |
| K15 | E7 | K15 | E7, F59 | K15 | D250, R3, T251 | K15 | R3, F59, D250, T251 |
| Q16 | F59 | Q16 | F59 | Q16 | F59, Y88 | Q16 | F59, Y60 |
| M17 |  | M17 |  | M17 | Y100 | M17 | Y100, I101 |
| A18 |  | A18 |  | A18 |  | A18 |  |
| V19 | F4 | V19 | F4 | V19 | F4, Y88, F56, F59 | V19 | F56, F4 |
| K20 | D90, Y88 | K20 | D90, Y88 | K20 | E92, D90, Y88 | K20 | E94 |
| K21 |  | K21 |  | K21 | E1 | K21 | E1, S163 |
| Y22 | N35, I8 | Y22 | N35, I8 | Y22 | I8, H5 | Y22 | I8, H5, L167 |
| L23 | Y88 | L23 | Y88 | L23 |  | L23 | Y88 |
| N24 |  | N24 |  | N24 |  | N24 |  |
| S25 |  | <u>**L25**</u> |  | S25 |  | <u>**L25**</u> |  |
| I26 |  | I26 |  | I26 | N35 | I26 | N35 |
| L27 |  | L27 |  | L27 |  | L27 |  |
| N28 |  | N28 |  | N28 |  | N28 |  |

## DISCUSSION

We created a library of novel VIP receptor antagonists based on cancer-associated mutations in the VIP gene and homologous sequences in the PHI peptide, estimated their docking scores by *in silico* modeling, synthesized antagonists with high predicted binding affinities to VPAC1 &VPAC2, and further evaluated their biological efficacies *in vitro and in vivo*. Our results showed that antagonists with better docking to VPAC1 & 2 effectively activate mouse T cells and enhance anti-leukemia response in murine models, including leukemia re-challenge experiments. ANT308 exhibits more than 10-fold lower EC_50_ to VIP-R on human T cells than VIPhyb (28); ANT308 significantly increased perforin-granzyme expression in AML patient-derived T cells and resulted in the highest survival rate in leukemia-bearing mice following daily subcutaneous administration. These findings align with our previous studies suggesting that VIP acts as an immune checkpoint molecule in various cancers, including AML, by inhibiting T cell activation (20, 23).

While earlier work demonstrated the potential of VIPhyb to modulate immune responses, its low potency and limited efficacy in preclinical models restricted its clinical applicability (30). Our novel VIP-R antagonists, particularly ANT308, can overcome the limitations of VIPhyb. The enhanced efficacy of ANT308 may be attributed to its predicted high binding affinity to both VPAC1 and VPAC2 (Fig. 1B), consistent with the hypothesis that more potent inhibition of VIP signaling leads to greater T cell activation and anti-tumor effects. Importantly, our study supports the emerging concept of VIP receptor signaling as a targetable pathway for cancer immunotherapy, a notion supported by the mutual exclusivity of VIP and PD-L1 expression in some cancers (20).

VIP C-termini modification gave rise to a variety of agonists to VIP receptors(34, 46); in contrast, the substitution of the N-termini sequence “HSDAVF” with “KPRRPY” from neurotensin (28), makes VIPhyb and novel molecules selected from our combinatorial library act as antagonists. To further understand the ligand-receptor interactions, we predicted and compared the binding mechanisms of VIP, ANT308 and VIPhyb to VPAC1 & VPAC2, respectively, by AlphaFold 3. Our simulation and ChimeraX predictions are consistent with cryogenic electron microscopy studies of VIP or PACAP27 to VPAC1 or VPAC2 (43, 44). The predictions of binding affinities to human VPAC1 are consistent between AlphaFold and *in silico* docking modeling performed by Creative BioLab with ANT308 _(highest affinity)_ > VIP _(medium affinity)_ > VIPhyb _(low affinity)_ (Table 5).

**Table 5.** Concordance of predicted binding affinities of peptides to human VPAC1 using AlphaFold and Creative Biolabs. AlphaFold predicted surrogate affinity scores (PAE_i) of ANT008, ANT300, ANT308, VIPhyb, VIP binding to VPAC1, with lower numbers representing higher predicted binding affinity. For comparison, binding affinities of the same peptides predicted by Creative Biolabs (Table 2) are show, with small, more negative numbers representing higher predicted binding affinity. ipTM is a modeling confidence score from AlphaFold that measures the accuracy of the predicted relative positions of the subunits forming the protein-protein complex. Values above 0.8 represent predictions with high precision, and values below 0.6 suggest a low probability of binding accuracy.

| Peptide | Binding affinity surrogate (PAE_i) w.r.t VPAC1 | Free energy of binding predicted by Creative Biolabs to VPAC1 | iPTM (modeling confidence) |
| --- | --- | --- | --- |
| ANT008 | 21.23 | -60.17 | 0.5 |
| ANT300 | 17.04 | -72.36 | 0.84 |
| ANT308 | 16.98 | -71.6 | 0.85 |
| VIPhyb | 21.67 | -60.62 | 0.48 |
| VIP | 18.7 | -65.8 | 0.88 |

We found that the N-terminal “KPRRPY” sequence predicted a β-turn in VIPhyb & ANT308 when bound to VPAC1 And VPAC2, limiting how deeply the peptides are inserted into the receptor binding pocket in the transmembrane region (Fig. 9). Next, we examined potential allosteric changes of intracellular domains of VPAC1 by binding to VIP or ANT308 (Fig.S1). However, no significant changes were noticed as superimposed docking portrayed minor deflections in VIP’s relative root mean square deviation (RMSD) compared to ANT308 binding to human VPAC1. However, subtle conformational change may be sufficient to open up or close downstream ligand binding sites and trigger or inhibit cytoplasmic signaling. Another possibility is that the low confidence (pLDDT<50) in the structural accuracy of the intracellular domain limited AlphaFold from correctly predicting subtle or even substantial structural alternations within intracellular region. Better virtual stimulation and structure-based rational design in the future would enhance the development of next-generation antagonists with improved clinical potential.

Despite the promising results, this study has several limitations. The screening strategy focused on first identifying peptides with high predicted binding affinities to VPAC1. Thus, the fact that these peptides have, in general, lower binding affinities to VPAC2 should not be interpreted as an intrinsic quality of all VIP receptor peptide ligands. Higher VPAC1 affinities (Fig. 1D, E) represent a selection bias related to the approach we used for peptide selection, and prioritizing VPAC2 binders would likely have yielded a different result. In addition, the basis for selecting specific residues for substitution into the VIPhyb sequence was based upon the identification of homologous sequences identified in VIP mutated in cancer or related peptides. Changing other residues could have a greater effect on binding affinity, but the very large potential universe of all peptide sequences (28^20^ permutations and combinations) makes an unsupervised algorithm for selecting peptides for screening impractical. The *in vivo* efficacy of VIP-ANTs was evaluated only in murine leukemia models, and it remains unclear whether the findings will translate to human patients with leukemia. Additionally, while we demonstrated enhanced T-cell activation *in vitro*, the variability in T-cell responses between different human donors (Figure 5) suggests that further studies are needed to fully understand the immunological effects of VIP-ANTs in diverse patient populations. The short half-lives predicted for peptides led us to treat mice with daily subcutaneous injections. While a single injection of ANT308 led to eliminating C1498 leukemia cells in some mice (data not shown), multiple daily injections were required to optimize anti-leukemia activity. Future studies are needed to explore whether longer-acting forms of the VIP-receptor antagonist might have greater efficacy or more convenient dosing schedules. Lastly, while the AlphaFold models provided useful structural predictions, experimental validation through cryo-electron microscopy or X-ray crystallography would be necessary to confirm the predicted receptor-antagonist interactions.

The translational implications of this study are significant. By demonstrating that more potent VIP receptor antagonists can enhance T cell-mediated anti-leukemia responses, this work suggests a potential new avenue for cancer immunotherapy targeting immune suppression in the tumor microenvironment. If these findings can be validated in human clinical trials, VIP-ANTs could complement existing immunotherapies, such as checkpoint inhibitors, particularly in cancers that overexpress VIP and are resistant to PD-1/PD-L1 blockade. Moreover, identifying ANT308 as a lead compound with favorable pharmacological properties supports its potential development as a therapeutic agent for AML and other cancers that exploit VIP signaling to evade immune surveillance.

## Materials & Methods

### Selection of novel VIP-ANT sequences

Novel VIP-ANT sequence design was based upon the combinatorial process of selecting VIP-related peptides substituted with the six N-terminal amino acid residues of neurotensin (as in VIPhyb, (28), 1 to 8 amino acid substitutions from the 8 VIP mutations identified in the Cancer Genome Atlas, as well as 4 amino acid residues of PHI that differ from VIP in aa positions 8, 9, 26, and 27. In addition, substitutions of alanine or histidine were made at each residue 1-28 of VIPhyb (alanine and histidine scanning,(35, 36), The target sequence of human VPAC2 protein (Uniprot ID: P41587) was obtained from the Uniprot database. The homologous template crystal structure of VPAC2 (PDB ID: 6NBF) was found by the BLAST program and further obtained by searching Protein Data Bank (http://www.rcsb.org/pdb). Next, the protein structure files were imported into Molecular Operating Environment (MOE), then the protonation status and hydrogen atoms positions were confirmed by employing LigX module. The environment condition was set to a temperature of 300K and a pH of 7. The homology modeling requires the alignment of protein sequences (VPAC2 versus Template) and the construction of 10 independent intermediate homologous models which are the results of the ranking selection of isomers with different loop structures and side chain flips. Finally, the homologous model with optimal score according to the Generalized Born/Volume Integral (GB/VI) scoring function (47)was selected and further energy minimization with AMBER10: EHT force field (51).

Creative Biolabs (Shirley, NY) performed two rounds of *in silico* screening for predicted binding affinity of the novel sequences to VIP receptors. First, 200 peptide sequences were screened for predicted binding to VPAC1, including the sequences of VIP, VIPhyb, and the various substitutions of 1-8 amino acid residues into the sequence of VIPhyb. Control peptides screened for VPAC1 binding included sequences with the randomly scrambled positions of amino acids 7-28 of VIP or VIPhyb or (leaving the N-terminal six amnio acids unchanged, VIP SCRAM and VIPhyb SCRAM). Second, 100 peptides with predicted high affinity to binding to VPAC1 and three control peptides were subsequently screened for binding to VPAC2. Based upon high predicted binding affinity to both human VPAC1 and VPAC2 (docking scores, kcal/mol), 12 selected peptide sequences were synthesized by RS Synthesis (Louisville, KY) and tested for their ability to increase proliferation of mouse and human T cells *in vitro* and inhibit the growth of leukemia in mice *in vivo* as detailed below.

### VPAC-VIP/VIPhyb/ANT308 complex structure model construction

The sequences for Human VPAC1 (PDB ID: 8e3z) and VPAC2 (PDB ID: 7vqx) were obtained through the Protein Data Bank (Europe). Three structure predictions were performed through the AlphaFold 3 server (AF) on the native ligand of VIP, the pre-existing antagonist, VIPhyb, and lead VIP-R antagonist candidate ANT308. As each run resulted in 5 model predictions, the one corresponding to the highest ipTM score was chosen to perform structural analysis using ChimeraX (48) The metric of pTM (predicted Template Modeling) provides a measure for accuracy for the overall structure of the complex by AF multimer, where a value of > 0.5 predicts congruence to true structures. However, ipTM (interface pTM) captures the predicted interface’s accuracy between the protein-protein complex subunits. This provides more confidence in finding accurate intermolecular bonds and interactions. All intermolecular hydrogen bonds and salt bridges (including atomic side-chain) between the binding ligand and receptor within a center-to-center atomic distance of 3.5 angstroms were found through structural analysis in ChimeraX. These provided us with all ligand-receptor bonds to find the most prominent residues actively involved in both the binding ligand and receptor chain by analyzing the frequencies of their interactions.

### PBMC, Cell lines and Mice

Healthy donor PBMC leukapheresis products were obtained from Stem Cell Technologies. De-identified PBMC from leukemia patients were provided by the Emory Winship Cancer Institute Cancer Tissue and Pathology Core. The C1498-myeloid leukemia cell line was obtained from ATCC. Dr. Bruce Blazar (University of Minnesota) provided the C1498-luciferase+ cell line C1498ff. Dr. Marcel van den Brink (City of Hope, Los Angeles) provided the P815 luciferase+ mastocytoma. DBA/2, C57BL/6 (CD45.2), B6 SJL (CD45.1), B6 albino (CD45.2), and B6 lucificerase+ mice (49), B6-L2G85, JAX stock #025854) were purchased from Jackson Laboratory (Bar Harbor, Maine). The mouse colonies were maintained at the Emory University Division of Animal Resources facilities. Both male and female mice were 8-10 weeks old.

### T cell Activation & Flow Cytometry

Human T cell Isolation was performed as described previously(50). In brief, PBMC cryo-samples were rinsed with PBS and rested overnight at 37°C in RPMI+10% FBS+50IU/ml IL-2. T cells were isolated from PBMC using human pan-T cell isolation kit, according to the manufacturer’s protocol (Miltenyi Biotec, Bergisch Gladbach, Germany, Catalog No. 130-096-535). Isolated T cells from 4–5 donors were pooled and seeded at a density of 1 × 10^6^/mL in 100 μL media in round-bottom wells in a 96-well plate, activated with a 1.5 μL/mL CD3/CD28/CD2 T cell activator (ImmunoCult, San Diego, CA, USA) in the presence of 50 IU interleukin 2 (IL2). Pooled T cells were activated in the presence or absence of VIP-ANTs and cultured for 48 h. Leukocyte Activation Cocktail with Golgi Plug (BD, Franklin Lakes, NJ, USA) was added 4 h prior to cell harvesting to assess CD69, Ki67, Granzyme B, Perforin expression in CD4+ and CD8+ T cells. Briefly, cells were stained with Fixable Aqua live/dead viability stain (1:100 dilution) for 5 min at room temperature (RT). Surface antibodies were added to the cells at the desired concentration and left to stain for 30 min at 4 °C. Following surface staining, cells were subsequently fixed and permeabilized for intracellular staining. Antibodies targeting Ki67, Granzyme B, Perforin were added and left to stain for 45 min at RT. List-mode files from stained samples were acquired a on five-laser Aurora cytometer (Cytek Biosciences, Inc, Fremont, CA) and analyzed using Flowjo software (Tree Star, Inc). For the murine T-cell proliferation assay, methods were modified from our previous work (38). MACS-column enriched splenic T-cells from luciferase+ B6 mice (B6-L2G85) were cultured at 100,000 cells per well in anti-CD3e-coated 96-well plates, with 30U/ml murine IL-2. Each VIP antagonist tested was added daily at 3 uM in quadruplicate wells for 3 days. After 72 hours of culture, T-cell proliferation was assessed by adding 30µg luciferin to quantify bio-luminescence using an IVIS Spectrum instrument and Living Image Software (PerkinElmer). Readings were normalized to represent % luminescence compare to control untreated wells.

### Mice leukemia cell injection and BLI image, and tumor measuring

All experimental procedures involving animals were approved by the Institutional Animal Care and Use Committee (IACUC) at Emory University. On day 0, B6 mice were injected by tail vein injection (i.v.) with 1 x 10^6^ C1498 cells. DBA/2 mice were injected subcutaneously (s.c.) with 1 x 10^5^ P815-luciferase mastocytoma cells. Growth of luciferase+ C1498ff in B6 albino recipients was assessed by bioluminescent imaging (BLI) after mice were anesthetized, injected intraperitoneally (i.p.) with luciferin substrate (15 μg/g mouse), and imaged using an IVIS imaging system, starting at day 6 before treatment and continuing once weekly. Albino B6 mice surviving to >60 days without evidence of AML (by luciferase assay) received a second injection of a two-fold higher dose of C1498 (2 × 10^6^ cells). DBA/2 mice that received P815 cells and treatment with ANT308, and survived past 60 days, were rechallenged using 2 x 10^5^ P815 cells. Lymphocyte kinetics and C1498 leukemia cell burden were analyzed weekly through pterygoid venous plexus blood collection in CD45.1+ B6.SJL recipients; CD45.2 C1498 tumor cells were differentiated from CD45.1 host cells by flow cytometry. Mice that reached endpoint conditions were euthanized, and counted as dead on the next day.

### VIP, VIPhyb, and VIP-ANT administration

Mice inoculated with acute myeloid leukemia cells were treated with subcutaneous injections of VIP, VIP scramble, VIPhyb, VIP-ANT [ANT008, ANT058, ANT107, ANT114, ANT195, ANT197, ANT203, ANT219, ANT300, ANT308], and control scrambled peptides [VIPhyb SCRAM, VIP SCRAM, Fully SCRAM, ANT308 SCRAM] peptides (RS Synthesis, Louisville, KY) (Fig. 1). A range of doses were administered subcutaneously at 3-30 nMoL/mouse. Treatment schedules tested included twice daily, once daily, once every other day, once every three days and once weekly, for 7-10 doses. Some mice received a single dose at day +6 following leukemia cell injection (Fig. 2A-C).

### CREB signaling & Western Blot

CREB signaling in T cells was measured as described previously (23). In brief, T cells from healthy donors pooled isolated and pooled, then cultured in complete RPMI containing 0.5% fetal bovine serum overnight. T cells were then incubated at 37°C in the presence of ANT308 at 10 μM for 30 min followed by stimulation with VIP for 15 min. T cells were then washed twice with ice-cold 1X PBS and lysed with ice-cold RIPA (R0278, Sigma) containing 1X protease inhibitor cocktail (P8340, Millipore Sigma) and phosphatase inhibitors (P2850, Millipore Sigma). Lysates were quantified by Bradford assay (BioRad), normalized for concentration, denatured with 1XSDS sample buffer. 40 μg of protein per sample was resolved by SDS-PAGE, blotted on PVDF membrane, and probed with primary antibodies. The images were acquired using BioRad ChemiDoc™ Touch Imaging System with Image Lab™ Touch Software.

### Cytospin, Immunofluorescence & Imaging

10^5^ P815 or C1498 cells were prepared in 100μl 1% BSA-PBS, Cytospin^TM^ centrifuged with Cytofunnels, Cytoslides and Cytoclips system (Thermo Scientific) for 3 min using medium acceleration. Cells on Cytoslides were then rinsed twice in PBS and fixed in 4% formaldehyde (Electron Microscopy Services, Hatfield, PA) at room temperature for 15mins, quenched with glycine, permeabilized with 0.3% Triton X-100 (Sigma), blocked with normal goat serum (Sigma). The cells were then incubated with primary antibodies in staining buffer overnight. After rinsing in staining buffer three times, the cells were incubated in secondary antibodies for 90 min. The cells were rinsed again three times, and mounted with Prolong^TM^ Gold Antifade Mountant (Invitrogen). Images cells in monolayer were captured with Leica SP8 Confocal Laser Point Scanning microscope with 63x/1.4 objective. Confocal Z-stacks were taken at 0.3 microns steps.

### Statistical analysis

Data were analyzed using Prism version 10, and SPSS statistics are presented as mean ± SD of all evaluable samples if not otherwise specified. Survival differences among groups were calculated with the Kaplan-Meier log-rank test pair-wise. Other data were compared using a one-way analysis of variance and nonparametric tests (Mann-Whitney U or Kruskal-Wallis H test). A p-value of < 0.05 was considered significant.

## Acknowledgements

Funding for this research was supported by a Georgia Research Alliance grant to Cambium Oncology and a Catalyst Drug Development award from the Winship Cancer Institute. Research reported in this publication was supported in part by the Cancer Tissue and Pathology (CTP) and Cancer Animal Models (CAMS) Shared Resources of Winship Cancer Institute Emory University and NIH/NCI under award number P30CA138292. The content is solely the responsibility of the authors and does not necessarily represent the official views of the National Institutes of Health.

