## Supporting Information for "Identification and characterization of vasoactive intestinal peptide receptor antagonists with high-affinity and potent anti-leukemia activity"

Supplemental Figure 1

A. VIP & VPAC1 B. ANT308 & VPAC1

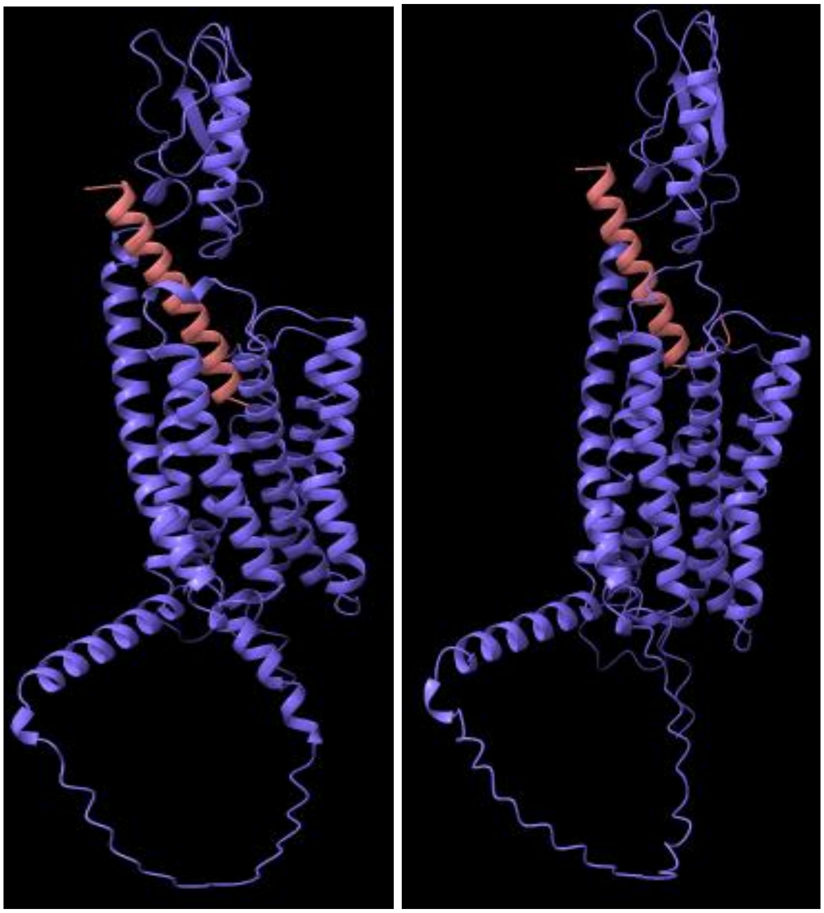

**Supplemental Figure 1**

**Human VIP-VPAC1, ANT308-VPAC1 complex solved by Alphafold.** Secondary structure comparison of human VPAC1 (purple multi-domain complex, full structure including transmembrane & intracellular domains) binding to VIP (pink peptide helices and beta sheets) or ANT308 (pink peptide helices and beta sheets).

Supplementary Figure 2

A. ANT300 & VPAC1 B. ANT008 & VPAC1

C. ANT300 & VPAC1 Zoom-In D. ANT008 & VPAC1 Zoom-In

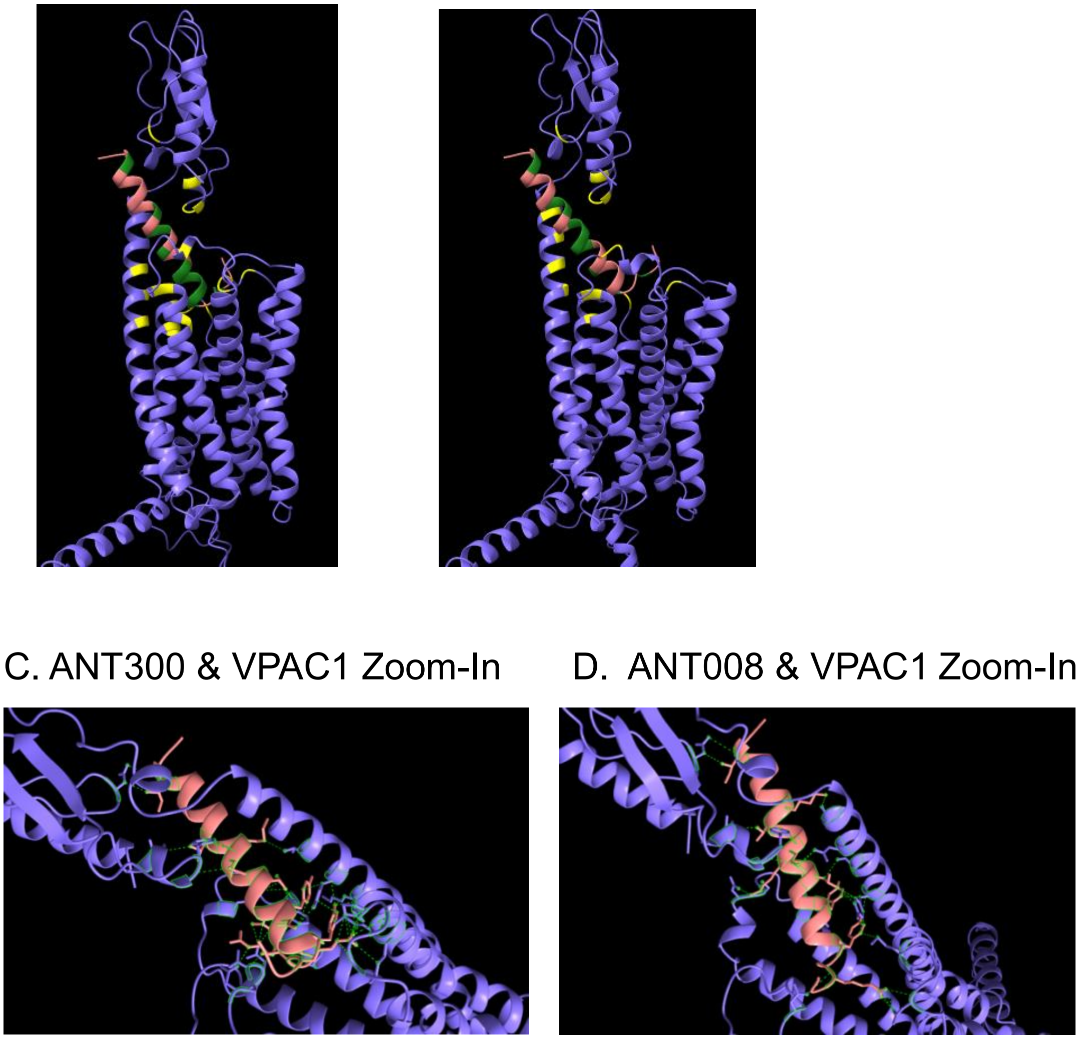

**Supplementary Figure 2**

**Human ANT300-VPAC1, ANT008-VPAC1 complex represented by Alphafold.**

Secondary structure comparison of human VPAC1 binding to ANT300, ANT008. Human VPAC1 (purple multi-domain complex, VIP binding interfaces highlighted in yellow). (*A)* ANT300, (*B)* ANT008 (pink peptide helices and beta sheets, VPAC1 binding interfaces highlighted in green). (*C-D)* Zoom-In view of ANT300 or ANT008 binding pockets at the transmembrane bundle of the VPAC1 receptor, pseudo bonds are highlighted in green.

Supplementary Figure 3

A. VIP & VIPR1

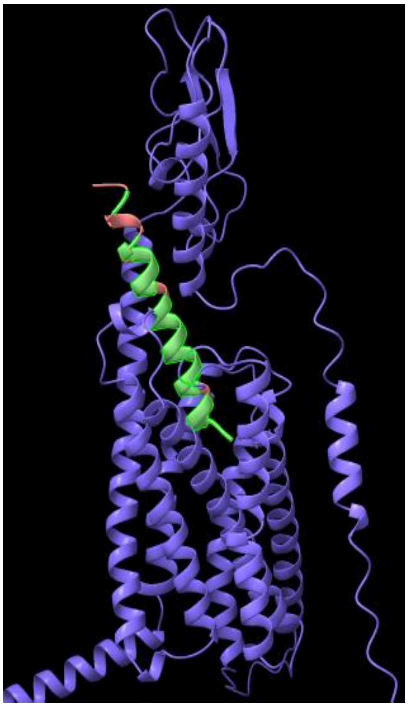

B. VIPhyb & VIPR1

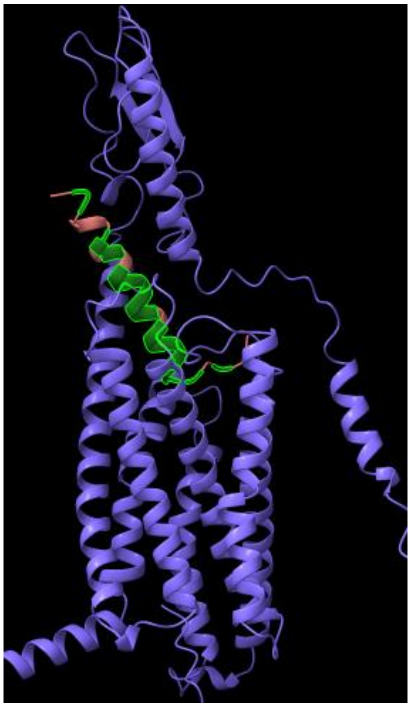

C. ANT308 & VIPR1

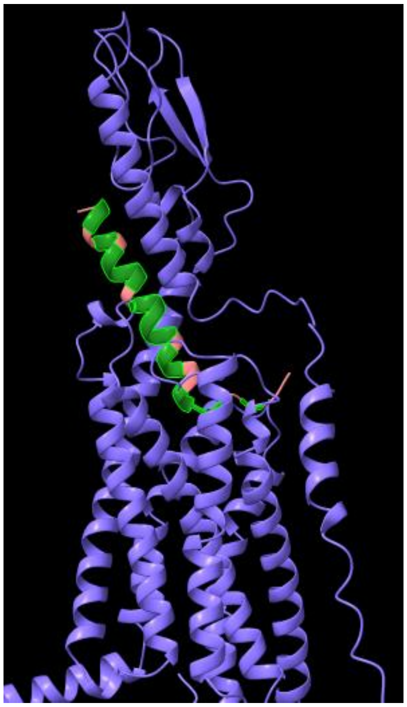

D. VIP & VIPR2

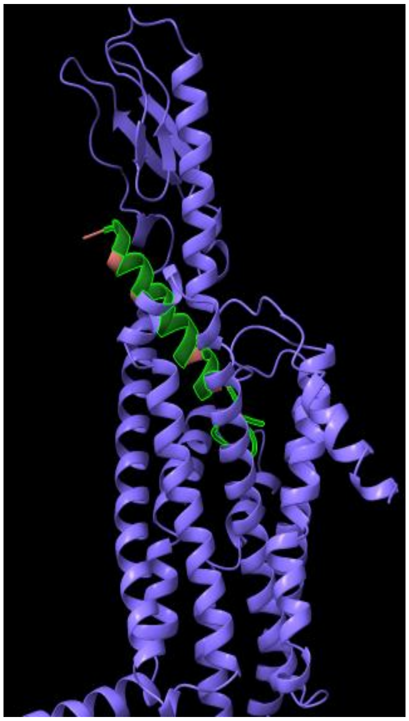

E. VIPhyb & VIPR2

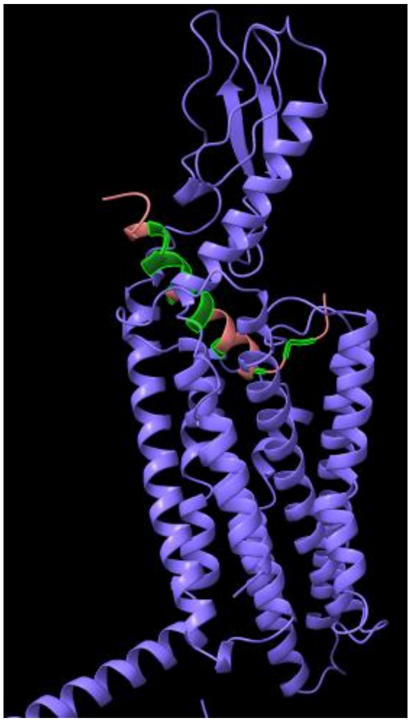

F. ANT308 & VIPR2

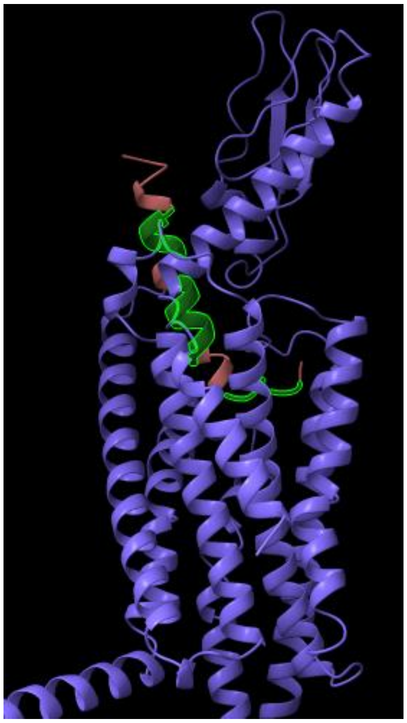

### Supplementary Figure 3

**The binding of peptide antagonists to mouse VIPR1, VIPR2 as represented by Alphafold.** Secondary structure comparison of mouse VIPR1 binding to VIP, ANT308, and VIPhyb. Mouse VIPR1 (purple multi-domain complex). *(A)* VIP, *(B)* VIPhyb, *(C)* ANT308 (pink peptide helices and beta sheets, VIPR1 binding interfaces highlighted in green). *(D-F)* Mouse VIP-VIPR2, VIPhyb-VIRP2, ANT308-VIPR2 complex solved by AlphaFold. Secondary structure comparison of mouse VIPR2 binding to VIP, VIPhyb, and ANT308. Mouse VIPR2 (purple multi-domain complex). *(D)* VIP, *(E)* VIPhyb, *(F)* ANT308 (pink peptide helices and beta sheets, VIPR2 binding interfaces highlighted in green).

Supplementary Table 1

| VIPhyb | Mouse VIPR1 | ANT308 | Mouse VIPR1 |
|---|---|---|---|
| ***<u>K1</u>*** | | ***<u>K1</u>*** | |
| ***<u>P2</u>*** | | ***<u>P2</u>*** | |
| ***<u>R3</u>*** | D364, K371 | ***<u>R3</u>*** | D364, K371 |
| ***<u>R4</u>*** | | ***<u>R4</u>*** | |
| ***<u>P5</u>*** | I290 | ***<u>P5</u>*** | I290 |
| ***<u>Y6</u>*** | Y140, M372, L376 | ***<u>Y6</u>*** | Y140, M372, L376 |
| T7 | K196 | T7 | K196 |
| D8 | I290, T289 | ***<u>S8</u>*** | |
| N9 | Y140 | ***<u>D9</u>*** | |
| Y10 | Y140, D141, F201, | Y10 | Y140, D141, F201 |
| T11 | D288 | T11 | D288 |
| R12 | | R12 | |
| L13 | Y140 | L13 | Y140 |
| R14 | L200, F201 | R14 | L200, H208, F201, N203 |
| K15 | E36 | K15 | P33 |
| Q16 | L92, F93 | Q16 | L92 |
| M17 | T137 | M17 | E133 |
| A18 | H208 | A18 | |
| V19 | | V19 | F93 |
| K20 | E133, F93 | K20 | E133, F93 |
| K21 | | K21 | E205 |
| Y22 | L40 | Y22 | L40 |
| L23 | | L23 | |
| N24 | | N24 | H119 |
| S25 | | ***<u>L25</u>*** | |
| I26 | N69 | I26 | N69 |
| L27 | Y118, L70 | L27 | Y118 |
| N28 | | N28 | |

**Supplemental Table 1. Amino acid residues involved in Mouse VIPR1 and peptide interactions predicted by ChimeraX.** Differences from VIP sequence are indicated by ***<u>bold, underlined, and italicized</u>*** letters. Residue numbers were counted directly within the Mouse VIRP1 sequence (UniProt, P97751).

Supplementary Table 2

| VIPhyb | Mouse VIPR2 | ANT308 | Mouse VIPR2 |
|---|---|---|---|
| ***<u>K1</u>*** | | ***<u>K1</u>*** | |
| ***<u>P2</u>*** | | ***<u>P2</u>*** | W280 |
| ***<u>R3</u>*** | P347 | ***<u>R3</u>*** | P347, I350 |
| ***<u>R4</u>*** | D275 | ***<u>R4</u>*** | D275 |
| ***<u>P5</u>*** | | ***<u>P5</u>*** | W280 |
| ***<u>Y6</u>*** | Q355, E359 | ***<u>Y6</u>*** | Q355, I356 |
| T7 | | T7 | |
| D8 | | ***<u>S8</u>*** | |
| N9 | | ***<u>D9</u>*** | |
| Y10 | Y129 | Y10 | K126, Y183 |
| T11 | | T11 | D272, Y183 |
| R12 | | R12 | N80 |
| L13 | | L13 | Y122 |
| R14 | D179, Y183, Y122 | R14 | Y183, K126 |
| K15 | D179, D272, T273, F81 | K15 | F81, R25 |
| Q16 | F81 | Q16 | F81, P113, Y110 |
| M17 | K118, I119, Y122 | M17 | |
| A18 | | A18 | |
| V19 | F78, F81, F26 | V19 | F26, F78 |
| K20 | D115 | K20 | D112, Y110 |
| K21 | E23 | K21 | E23 |
| Y22 | I30, H27 | Y22 | I30, N57, F26, H27 |
| L23 | Y110, F78 | L23 | |
| N24 | | N24 | |
| S25 | | ***<u>L25</u>*** | |
| I26 | | I26 | |
| L27 | | L27 | |
| N28 | | N28 | |

**Supplemental Table 2. Amino acid residues involved in Mouse VIPR2 and peptide interactions predicted by ChimeraX.** Differences from VIP sequence are indicated by ***<u>bold, underlined, and italicized</u>*** letters. Residue numbers were counted directly within the Mouse VIRP2 sequence (UniProt, P41588).

### Antibodies for Western Blots

| Target | Origin | Reference | Catalog # | Concentration |
|---|---|---|---|---|
| Phospho-CREB (Ser133) | Rabbit monoclonal | Cell Signaling Technology | 9198 | 1 : 1000 |
| CREB | Rabbit monoclonal | Cell Signaling Technology | 9197 | 1 : 1000 |
| β-actin | Mouse monoclonal | Cell Signaling Technology | 3700 | 1 : 1000 |

### Antibodies for Flow Cytometry

| Target | Clone | Fluorochrome | Supplier | Reactivity | Catalog# | Dilution |
|---|---|---|---|---|---|---|
| CD3 | UCHT1 | PE-CF594 | BD Biosciences | Human | 562310 | 1 to 100 |
| CD8 | SK1 | FITC | BioLegend | Human | 344704 | 1 to 100 |
| CD4 | RPA-T4 | APC-Cy7 | BioLegend | Human | 300518 | 1 to 100 |
| CD69 | FN50 | PE-Cy7 | BioLegend | Human | 310912 | 1 to 100 |
| Perforin | dG9 | Alexa Fluor 647 | BioLegend | Human | 308110 | 1 to 100 |
| Granzyme B | GB11 | Pacific Blue | BioLegend | Human | 515408 | 1 to 100 |
| Ki67 | Ki-67 | BV605 | BioLegend | Human | 350522 | 1 to 100 |
| VIP | OT15B5 | PE | OriGene | Human, Mouse | CF806852 | 1:50 |
| Mouse IgG1 | MOPC-21 | PE | BD Parmingen | Mouse | 555749 | 1 to 100 |
